# Robust Evaluation of Deep Learning-based Representation Methods for Survival and Gene Essentiality Prediction on Bulk RNA-seq Data

**DOI:** 10.1101/2024.01.23.576852

**Authors:** Baptiste Gross, Antonin Dauvin, Vincent Cabeli, Virgilio Kmetzsch, Jean El Khoury, Gaëtan Dissez, Khalil Ouardini, Simon Grouard, Alec Davi, Regis Loeb, Christian Esposito, Louis Hulot, Ridouane Ghermi, Michael Blum, Yannis Darhi, Eric Y. Durand, Alberto Romagnoni

## Abstract

Deep learning (DL) has shown potential to provide powerful representations of bulk RNA-seq data in cancer research. However, there is no consensus regarding the impact of design choices of DL approaches on the performance of the learned representation, including the model architecture, the training methodology and the various hyperparameters. To address this problem, we evaluate the performance of various design choices of DL representation learning methods using TCGA and DepMap pan-cancer datasets, and assess their predictive power for survival and gene essentiality predictions. We demonstrate that non DL-based baseline methods achieve comparable or superior performance compared to more complex models on survival predictions tasks. DL representation methods, however, are the most efficient to predict the gene essentiality of cell lines. We show that auto-encoders (AE) are consistently improved by techniques such as masking and multi-head training. Our results suggest that the impact of DL representations and of pre-training are highly task- and architecture-dependent, highlighting the need for adopting rigorous evaluation guidelines. These guidelines for robust evaluation are implemented in a pipeline made available to the research community.

## Introduction

Precision medicine and the development of new therapies require accurate disease diagnosis and outcome prediction. The field of omics research has experienced an unprecedented data revolution fueled by high-throughput technologies, enabling the generation of high-dimensional omics data at an exponential pace. This wealth of data provides interesting opportunities to unravel the molecular landscape of diseases, including cancer, and emphasizes the need for robust computational approaches to extract meaningful insights. In particular, RNA sequencing (RNA-seq) is now ubiquitous in molecular biology and oncology (Stark, Grzelak, et Hadfield 2019) and was shown to be the most informative omics modality for predicting phenotypes of interest such as patient survival (Vale-Silva et Rohr 2021) or gene essentiality in cell lines (Chiu et al. 2021).

In parallel, deep learning-based representation learning approaches have shown remarkable potential in analyzing complex data, ranging from images to text (Bengio, Courville, et Vincent 2013; Devlin et al. 2019; Misra et Van Der Maaten 2020). These methods, powered by artificial neural networks, excel at capturing intricate patterns, detecting subtle relationships, and making accurate predictions. Applying deep representation learning techniques (DRL) to RNA-seq data for cancer research holds the potential to revolutionize our understanding of cancer progression, classification, and treatment response.

Therefore, the integration of deep learning-based approaches within the field of omics research holds immense promise for advancing our understanding of cancer biology (Chaudhary et al. 2018). Nonetheless, despite the vast potential of DRL algorithms and demonstrated success in vision and Natural Language Processing (NLP) domains, they still face challenges in surpassing traditional tree-based methods on tabular data (Grinsztajn, Oyallon, et Varoquaux 2022). Importantly, their application to omics data remains underexplored when considering gene expression matrices derived from bulk RNA-seq.

Typical tasks associated with omics data like survival or gene essentiality predictions present unique challenges, involving high-dimensional feature spaces with limited sample sizes, a need to account for batch effects and variations in data generation procedures (J. Liu et al. 2018; Gönen et al. 2017). They require various intricate steps like normalization, scaling, dimensionality reduction (DR), and the selection of prediction models and training frameworks. Moreover, the presence of noisy and heterogenous labels, such as survival and censoring information, further complicates the analysis. Generally, these tasks can be sensitive to overfitting and exhibit significant variability across different datasets and tasks.

Reducing the dimensions of genomic data is often a privileged option to help address such difficulties. Non-DL dimensionality reduction methods have proven effective in analyzing omics data for guided feature selection (Zhakparov et al. 2023), deconvolution of bulk RNA-seq data (Y. Liu et al. 2018), clustering (R. Chen et al. 2022; Wei et al. 2023) or prediction of clinical endpoints (Sauta et al. 2023). With the rise of deep learning, several papers have explored new representation learning methods for cancer transcriptome data analysis such as auto-encoder architecture variations for biomarker identification (Q. Li et al. 2023; De Weerd et al. 2023), subtyping (Withnell et al. 2021) and supervised tasks including gene essentiality (Chiu et al. 2021) or drug response predictions (He et al. 2022; J. Chen et al. 2022; Dincer et al. 2018; Rampášek et al. 2019). In single-cell data, the higher amount of data points allowed the development of more complex methods inspired from self-supervised learning (Shen et al. 2021; Han et al. 2021), graph-based methods (X. Li et al. 2017) and more recently from large language models (Theodoris et al. 2023; Cui et al. 2023; Shen et al. 2023). Nonetheless, despite numerous publications showcasing the potential of deep representation learning, there is a scarcity of comprehensive benchmarks in the context of cancer research focusing on deep-learning methods for bulk RNA-seq due the multitude of potential tasks. Existing benchmarks showcase limited improvement over linear baselines when using deep learning methods in phenotype prediction or clustering (Smith et al. 2020; Cantini et al. 2021), but do not cover tasks such as gene essentiality prediction or pan-cancer pretraining approaches. This lack of reliable comparisons seemingly originates from the difficulty of accurately assessing models’ performances in a simple cross-validation setting (Bengio et Grandvalet 2003; Nadeau et Bengio 2003), which becomes even more complex when adding hyperparameters (HPs) tuning, small dataset sizes, noisy labels and confounding factors, which are ubiquitous in omics for oncology (Whalen et al. 2021).

This impedes the development and evaluation of these methods, limiting their reliability and usage. It has been argued that the rapid growth of deep learning in other applications can be partly attributed to the widespread adoption of clear benchmarks, like ImageNet (Deng et al. 2009) for visual deep learning or more recently the CASP competition for protein structure prediction (Kryshtafovych et al. 2021).

In this paper, we investigated and robustly benchmarked the performance of different prediction models and training approaches on bulk RNA-seq data in the context of cancer research. In particular, we studied the impact of the representation method choice on the performance of 11 cancer-specific survival prediction tasks, a pan-cancer survival prediction task on 33 combined cohorts and a gene essentiality prediction task on hundreds of cell lines. We implemented DRL architectures that reach state-of-art performances in other fields, namely auto-encoders (AE), variational auto-encoders (VAE), masking auto-encoders (MAE), multi-head auto-encoders (MHAE), data augmentation techniques coupled with AE (DA), graph neural network (GNN) based on prior knowledge approaches, pre-trained models (PreAE), and compared them with standard baseline representations (Identity and PCA). Our evaluation encompassed preprocessing steps such as normalization, scaling, and feature selection, and provided a fully reproducible framework to evaluate their impact as well as the one of random splits during cross-validation and hyperparameters (HPs) tuning strategy on different prediction tasks.

This study highlights the considerable variability in results due to data splits and the limited impact of the choice of the representation model or training framework on the performances on the considered tasks.

## Materials and Methods

In this section, we describe all the different elements of the benchmarking pipeline depicted in **Supp. Fig. S1.** Additional details can be found in Supplementary Materials. Although our pipeline can be used to evaluate any RNA-seq-based ML model, we focused in this study on benchmarking the capacity of various representation models to capture relevant information in bulk RNA-seq data.

### Datasets

This study was based on publicly available data from The Cancer Genome Atlas Program (TCGA) and Cancer Cell Line Encyclopedia (CCLE) dataset (Barretina et al. 2012). Expression data (raw counts, RPKM, TPM) for all TCGA datasets were downloaded from RECOUNT3 (Wilks et al. 2021), a publicly available resource concatenating multiple omics datasets aiming at harmonizing preprocessing steps to make comparisons easier between these datasets. For the per-cohort Overall Survival (OS) prediction task, we focused on the cohorts with less than 90% of patients censored and selected the indications with more than 400 patients with associated RNA-seq samples (BRCA, UCEC, KIRC, HNSC, LUAD, LGG, LUSC, SKCM, COAD, STAD and BLCA). For the pan-cancer task, we selected the 33 cohorts of TCGA, for a total of 10,736 samples. For both OS prediction tasks, we used the overall survival (OS) labels defined by the TCGA Pan-Cancer Clinical Data Resource (TCGA-CDR) (J. Liu et al. 2018).

The Gene Essentiality prediction experiments were carried out using the CCLE dataset, which contains the expression profiles of hundreds of human cancer cell lines. The dataset was obtained through the DepMap portal, version 22Q4 (Barretina et al. 2012), which provided the gene perturbation experiments results used as labels. Expression data (raw and TPM pseudo-counts) and gene signatures files were downloaded directly from the platform and non-transformed TPM data was obtained by applying a reverse transformation of the pseudo-counts file. We used the same selection process as previous studies (Chiu et al. 2021) to define gene dependencies of interest and filtered out cell lines with missing dependencies. This resulted in 1,223 genes of interest and 893 cell lines for our experiments.

### Tasks description

The different representation architectures were evaluated by the performance on the prediction of patients or cell lines labels when using the output of baseline representation models or of DRL ones as input features. These downstream tasks were evaluated on several test sets sampled from the datasets following a repeated holdout cross-validation process (see related section below for more details).

The first task we considered was cancer-specific overall survival (OS) prediction, a common task in cancer biology that can improve the identification of subgroups of patients at risk. In this task, representation and prediction models were trained on single cancer cohorts separately to predict the survival of patients within a particular cancer subtype. Our second task consisted in pan-cancer OS prediction, which is often considered in the literature as a standard for comparing methods and claim state of the art performance of newly developed ML models (Vale-Silva et Rohr 2021). The models are trained on all the cohorts from TCGA combined in a single dataset, evaluating the capabilities to predict the different survival times of a more heterogeneous population suffering from different indications. Harrell C-index (Harrell, Lee, et Mark 1996) was used as the final metric for both tasks and is referred to as c-index in the paper. The different data splits were stratified to ensure similar censorship levels between train and test sets.

The gene essentiality prediction task described in this paper was inspired by the DeepDEP framework (Chiu et al. 2021), with the modification of predicting gene dependency instead of gene effect as recommended in (Dempster et al. 2019; Chiu et al. 2021) and focusing only on bulk RNA-seq rather than a multi-modal setting. In this context, gene dependency refers to the extent to which a particular gene is necessary for cell survival and growth, which depends on which cell line is considered. Each cell line is represented by its expression data while genes are represented by their fingerprints. Fingerprints are specific encodings that summarize the relevant biological functions of a given gene and represent its involvement in 3,115 gene signatures related to chemical and genetic perturbation defined in MSigDBv6.2 (Liberzon et al. 2011). The inputs for representation models were the RNA-seq expression matrix and the gene fingerprints, while the output was then passed to the prediction models. We focused only on the cell line representation based on RNA-seq data in our benchmark while the fingerprints were fixed and reduced to 500 dimensions using PCA, an arbitrary choice to balance between computation speed, performance and variance explained (**Supp Figure S2**). For each cell line / gene combination with a gene dependency label, we created the corresponding features by concatenating the cell line representation and the fingerprints representation. To evaluate model performances, we used the Spearman rank correlation. While in the original DeepDEP paper Pearson correlation coefficients were computed, both methods are suited to evaluate this task (Rosenski, Shifman, et Kaplan 2023; Ma et al. 2021). The Spearman correlation was preferred as it is more robust to outliers on non-normally distributed data (Hou et al. 2022) and fits better the framework of ranking genes for target selection. The correlation was computed on the whole test set and is referred to as the overall correlation. Following DeepDEP’s evaluation, we computed as well a per-gene metric: for each gene with available experiments, we calculated the Spearman rank correlation over the different cell lines and averaged the result over all the genes to have one final metric per repetition. To ensure a rigorous evaluation we partitioned the data by cell lines, ensuring that a cell line did not overlap between the training and test sets.

### Preprocessing Datasets and Gene Selection

The RNA-Seq data was preprocessed following a classical bioinformatics pipeline. While different preprocessing options were tested (data and results not shown) as they can impact downstream analyses (Paton et al. 2023), we decided to choose a fixed standard choice to keep a reasonable computational budget and compare models all else being equal. For all the experiments without pre-training on external datasets, we selected the 5,000 most variable genes on the training sets, applied a logarithmic operation and normalized the data with mean-standard scaling. The exact same transformation was then applied to the test dataset to prevent any potential leak by computing features statistics on the whole dataset. All models used TPM normalized expression data except the variational auto-encoder model model which used raw counts to fit a negative binomial prior distribution (*cf Methods, Representation Models*).

When considering pre-training, we implemented another preprocessing process closer to recent pre-training strategies for foundation models (FM) trained on single-cell expression data (scRNA-seq) (Cui et al. 2023; Shen et al. 2023; Theodoris et al. 2023). While these models can afford little feature selection during pre-training thanks to the vast amount of available data, pre-training on size-limited bulk RNA-seq datasets still requires gene selection to avoid the curse of dimensionality. However, the most variable genes for the pre-training datasets can be different from those of the datasets used to evaluate on the downstream tasks for non-pretrained models. To select genes that were still relevant for the tasks considered in this paper, we built two gene lists based on their variances in the TCGA pan-cancer dataset used for pretraining :

● In the case of the per-cohort OS prediction task, the previous procedure has to be adapted in order to ensure a fair comparison with the non-pre-trained case. We therefore took the union of the top 2,000 most variant genes for each of the 11 cohorts selected in the downstream task to make sure relevant genes per indication were also selected in the final features rather than genes solely linked to cancer type that would be considered when looking at the genes variances across the whole TCGA dataset. This resulted in a set of 5,046 unique gene identifiers.
● For the gene essentiality task, we took the top 5,000 most variant genes in TCGA after intersection with the genes present within the CCLE dataset, similarly to DeepDEP’s procedure of selecting genes with a standard deviation superior to 1 in TCGA.

The TCGA data used for pretraining was normalized within each fold with mean standard scaling and learned statistics were saved for potential usage on the downstream datasets (Pre-training experiments).

### Repeated holdout cross-validation framework

In this study, we aim to compare the performance of different representation learning algorithms on downstream survival and gene essentiality prediction tasks using bulk RNA-seq data. Each representation model is trained and used to transform the input expression data before feeding the learned low-dimensional embeddings to a task-specific prediction model, fitted for each representation model tested. To achieve a comprehensive evaluation, we adopt a rigorous validation pipeline that focuses on exploring the learning algorithm’s variability to diverse hyperparameter settings. Our validation pipeline involves a repeated holdout cross-validation approach (Varoquaux et Colliot 2023) in which the dataset is repeatedly split in two to create pairs of training and test sets, comprising 80% and 20% of the original data respectively (**Supp Fig S1**). For experiments without pre-training, the training sets are used to select jointly the optimal HPs for the representation and prediction models by performing a 5-fold cross-validation for a given set of HPs. The HP tuning is performed using a Tree-structured Parzen Estimator (TPE Sampler) implemented by Optuna (Akiba et al. 2019) with a fixed budget of 50 iterations. Then, we select the set of HPs with the best average performance over the validation folds on the downstream tasks to train the representation and prediction models on the whole training set before evaluating it on the test set. This procedure is repeated 10 times to generate a distribution of scores over the different test sets, providing robust performance assessments compared to a single test evaluation.

For pre-training experiments, we performed for each task HP tuning of the pre-trained representation models separately from the prediction models. We used the TPE Sampler with 50 trials to find the architecture choices and regularization factors (**Table S1**) that minimized the reconstruction loss of the auto-encoders on TCGA data. For each trial, a 5-fold cross-validation was repeated 5 times to estimate the generalization error by averaging the scores obtained per fold. For the gene essentiality prediction task, all of the 33 cohorts were used during pre-training (for a total of 10,736 samples in the pretraining dataset) while we removed the 11 cohorts of the downstream tasks for the per-cohort OS prediction task (4,480 samples in the pretraining dataset).

### Acceptance Testing and Model Scoring

In (Varoquaux et Colliot 2023), the authors critically examine the limitations of traditional null-hypothesis significance testing and the cross-validation framework in deriving statistical conclusions. They highlight the unsuitability of standard null-hypothesis significance testing, including t-tests, for cross-validation due to the non-independence of runs and the complexity of deriving confidence intervals. Consequently, they advocate for a repeated holdout framework as an alternative to cross-validation.

While it is feasible to derive confidence intervals from the repeated holdout framework, the authors still recommend a different scoring system. They argue that traditional hypothesis testing primarily focuses on the statistical significance of expected improvements in models over an infinite population. This approach, however, is not applicable to studies that concentrate on practical, meaningful improvements on finite-sized test sets. Therefore, following the authors’ recommendation, we considered a method superior if it outperforms another 75% of the time.

We proposed a scoring system for all models considered for a given task based on the aforementioned criterion. To evaluate the relative importance of each model, a score was generated by counting the number of significant pairwise comparisons won against other models. When different cohorts were considered for the same task, an average score was obtained from the cohort-specific scores. This value was then normalized by the maximum value obtained by a model on the task, resulting in a score between 0 and 1. This score was then converted into a percentage to rank the different models.

### Representation Models

We selected state-of-the-art methods from various subfields of DRL: linear models as baselines (Identity, PCA), auto-encoders-like models (AE, VAE), Self-Supervised Learning methods (Masking Auto-Encoders), Semi-Supervised methods (Multi-Head Auto-Encoders), data augmentation techniques coupled with AE, graph-based methods leveraging prior-knowledge (GNN) and training frameworks (Pre-training). For each of these categories, one base model was implemented and we considered the different architecture choices as HPs sweeps in our pipeline as described above. Details about the architectures and HPs ranges (Supp. Table S1) are described in Supplementary Materials.

#### Baselines

We considered as baseline representations the Identity and Principal Component Analysis (PCA), widely used for all considered tasks on RNA-seq data. The number of components of the PCA was considered a hyperparameter and optimized per fold in our pipeline (Supp. Table S1).

#### Auto-Encoders

An auto-encoder (AE) is a type of neural network that learns to encode input data into a lower-dimensional representation and then decode it back to the original form, with the goal of reconstructing the input accurately (Hinton et Salakhutdinov 2006). Given the limited size of our datasets, we focused on small architectures ranging from 0 to 2 hidden layers for the encoder (excluding the representation layer) with 256 to 1024 neurons (more details available in Supp. Table S1). The decoder part was constructed symmetrically to the encoder. We optimized the AE models using a Mean Squared Error (MSE) Loss and the Adam algorithm (Kingma et Ba 2017) and applied rectified linear unit activation (ReLU) functions between each linear layer. We did the same for other representation models derived from this architecture (MAE, MHAE, DA-GN, PreAE).

#### Variational Auto-Encoders: scVI

A variational auto-encoder (VAE) is a type of auto-encoder that incorporates a probabilistic approach, using encoder and decoder networks to generate latent variables and enable sampling from a learned distribution, allowing for generative modeling and capturing underlying data distributions (Kingma et Welling 2013).

Specifically, we adapted a popular method to embed scRNA-seq datasets, scVI (Lopez et al. 2018) to bulk RNA-seq data, and assessed the learned representations on our benchmark tasks. scVI is based on a hierarchical Bayesian model where conditional distributions are parametrized by neural networks. As in the VAE, each gene expression profile is encoded through a non linear transformation into a low dimensional latent vector. This latent representation is then decoded by another non linear transformation to generate an estimate of the parameters of a Zero-Inflated Negative Binomial. Since bulk RNA-seq data typically does not fit the zero-inflation assumption observed in scRNA-seq, we parameterized scVI’s decoder with a Negative Binomial distribution instead.

#### Masking Auto-encoders

Masking in self-supervised learning refers to the process of randomly hiding or removing portions of input data, forcing the model to learn to reconstruct the missing parts, which promotes the discovery of meaningful features and representations. VIME, or Value Imputation and Mask Estimation is a popular masking method for tabular data (Yoon et al. 2020). In this self-supervised learning framework for tabular data, two pretext tasks are introduced: feature vector estimation and mask vector estimation. The encoder function is trained together with two pretext predictive models to reconstruct a corrupted input sample (feature vector estimation) with MSE loss and estimate the mask vector used to corrupt the sample (mask vector estimation) with a cross-entropy loss. The resulting learned representation captures correlations across different parts of the data, making it useful for downstream tasks. We focused on one re-implementation from the main paper which is similar to the original method where we only included one pretext task: feature vector estimation.

#### Multi-head Auto-Encoders

A multi-head auto-encoder (MHAE) includes one or more auxiliary heads to perform supervised tasks during the representation model training. These auxiliary heads are fully-connected neural networks that take the compressed representation as inputs, and predict labels such as overall survival and gene essentiality as outputs. In this study we considered MH auto-encoders trained with an auxiliary head predicting the label of the downstream task, to assess if adding supervision would improve the performance of the auto-encoder.

#### Graphs Neural Networks

We used a Graph Neural Network (GNN) architecture to incorporate protein-protein interaction networks (PPIs) as a source of prior knowledge. The STRING PPI database served as the underlying graph on which the bulk RNA-seq data was laid. Each node of the graph represented a gene, with gene expression as a node feature. In this setting, each patient (or sample) was represented as a single graph, and the graph topology over the samples did not vary, only the overlaying signal did. This model is close to the traditional convolutional neural network alternating between convolution and pooling steps, but using a graph instead of e.g. pixel coordinates to perform message passing to neighboring features. (Althubaiti et al. 2021; Ramirez et al. 2021) previously showcased this kind of model for survival prediction, and we propose here a modified version intended for representation learning. More details about the implementation are given in the Supplementary Materials.

#### Data Augmentation

Data augmentation is a set of techniques used to reduce overfitting of machine learning models by generating new training data points from the original ones. It is ubiquitous in computer vision, where new images can be obtained by simple transformations such as rotations or cropping (Perez et Wang 2017). In the case of omics data, there are no obvious equivalents of such transformations. However, certain methods have been successfully applied to single-cell RNA-seq data in the context of contrastive learning by adding noise and simulating dropout events (Han et al. 2021). Specifically, we focused on using data augmentation based on Gaussian Noise (DA GN). The method consists in creating additional samples by copying the data and adding gaussian noise to all the features after they have been standardized with mean and standard normalization. We used a centered normal distribution with standard deviation controlled by a hyperparameter and fixed the number of copies to 4 corrupted samples for one original sample. The target labels for these new data points are simply copied from the original dataset.

### Pre-training experiments

The use of bulk RNA-seq data to train representation algorithms is often hindered by the limited number of patients who were screened for a specific condition. To address this issue, pre-training models on larger datasets can help represent the data more accurately. In this study, we tried two different pre-training strategies for both the per-cohort survival prediction and the gene essentiality prediction tasks. In order to better assess the effect of pre-training and decouple it from architecture details, we focused our pre-training experiments on the AE model.

For both downstream tasks, we compared the original AE to two different pre-training strategies, **PreAE** and **PreAE finetuned.** In the case of PreAE, the mean-standard scaling was done using solely the statistics of the TCGA pre-training dataset (as described above, all 33 TCGA cohorts were used for pre-training for gene essentiality prediction task, while we removed the 11 cohorts of the downstream tasks for the per-cohort OS prediction task) and the auto-encoder was used only for inference on the task dataset. For the PreAE finetuned strategy, the scaling was performed using the learned statistics on the training folds of the task dataset while the auto-encoder was also fine-tuned for each fold. These former experiments assess how initializing the weights with pre-training can help on the downstream tasks. Both optimization histories for the pretraining are available in (**Supp Fig S3, Supp Fig S4**).

### Downstream prediction models

For survival prediction tasks, each representation model was combined with a multi-layer perceptron (MLP) optimized with a differentiable Cox loss (Faraggi et Simon 1995; Katzman et al. 2018) using the Adam algorithm. 20% within each training fold were used for early stopping to prevent overfitting of the MLP. The choice of using MLP rather than Cox linear models was motivated by faster computation times thanks to GPU usage with PyTorch and by the need of a fair comparison with DL models that use nonlinear prediction heads. More details about the hyperparameters of the prediction model are available in the Supplementary Materials (**Table S2**).

For Gene Essentiality a light gradient-boosting machine (LGBM) regressor was trained to predict the essentiality (dependency score) of each gene in a cell line based on the features described in the task section. To train the LGBM regressors effectively, two HPs were considered: the learning rate and the regularization coefficient alpha, the other HPs having minimal impact in our experiments (data not shown). The learning rate was set in the range of 0.01 to 0.3 and the regularization coefficient within a range of 0 to 100 for exploring various regularization strengths. Notice that we initially tested both MLP (the choice in DeepDEP paper) and LGBM but ended up keeping the best performing model (data not shown).

## Results

In this work, we benchmarked the performance of different representations of bulk RNA-Seq data on survival prediction tasks on TCGA dataset and gene essentiality prediction on DepMap data. In particular, we compared the results obtained with standard baseline representations (Identity and PCA) with those obtained with DRL architecture such as auto-encoders (AE), variational auto-encoders (VAE), masking auto-encoders (MAE), multi-head auto-encoders (MHAE), data augmentation techniques coupled with AE (DA), graph neural network (GNN) based on prior knowledge approaches, and pre-trained models (PreAE) (see Materials and Methods).

The benchmarking pipeline (Supp. Fig. S1) was based on repeated holdout cross-validation processes and addressing issues such as unfair budget for HP tuning, overfitting or performing hardly generalizable statistical tests on particular data splits (See Materials and Methods section and Supplementary material section).

### Comparing DRL and baseline models performances

We first considered the survival prediction tasks in TCGA datasets by studying separately the different cancer cohorts. This task is known to be challenging, as the OS labels are noisy (high percentage of censoring and debatable quality of the labels as shown in (J. Liu et al. 2018)), challenging to predict accurately, and with high variability of model performance across different cohorts (Huang et al. 2020). As shown in **Fig 1**, in the indication-specific survival prediction task, we observed that the choice of the representation model had minimal impact on performance, as they consistently achieved similar scores in terms of mean c-index, with less than 10% difference in most cohorts between the best and worst models and close to 15% difference in BRCA or STAD. Interestingly, Identity and PCA demonstrated excellent performance on most cohorts. Notably, as shown in **Fig 1** and **Supp. Fig S5**, using 75% acceptance criterion on test folds, no model outperformed all the others in at least one cohort. Nonetheless, by using the number of significant pairwise comparisons, Identity was the best representation model choice for BRCA (together with MAE), LUAD, COAD (together with AE) and BLCA while PCA was the best choice for UCEC, KIRK and HNSC. Among the DRL methods, under the same criteria, AE is also the best choice for SKMC and STAD, MHAE for LGG. Notice that these results are not equivalent to standard comparison of mean performance (cfr. Supp. Table S3), for which for example Identity would only be considered the best model for HNSC and BLCA.

**Figure 1.**
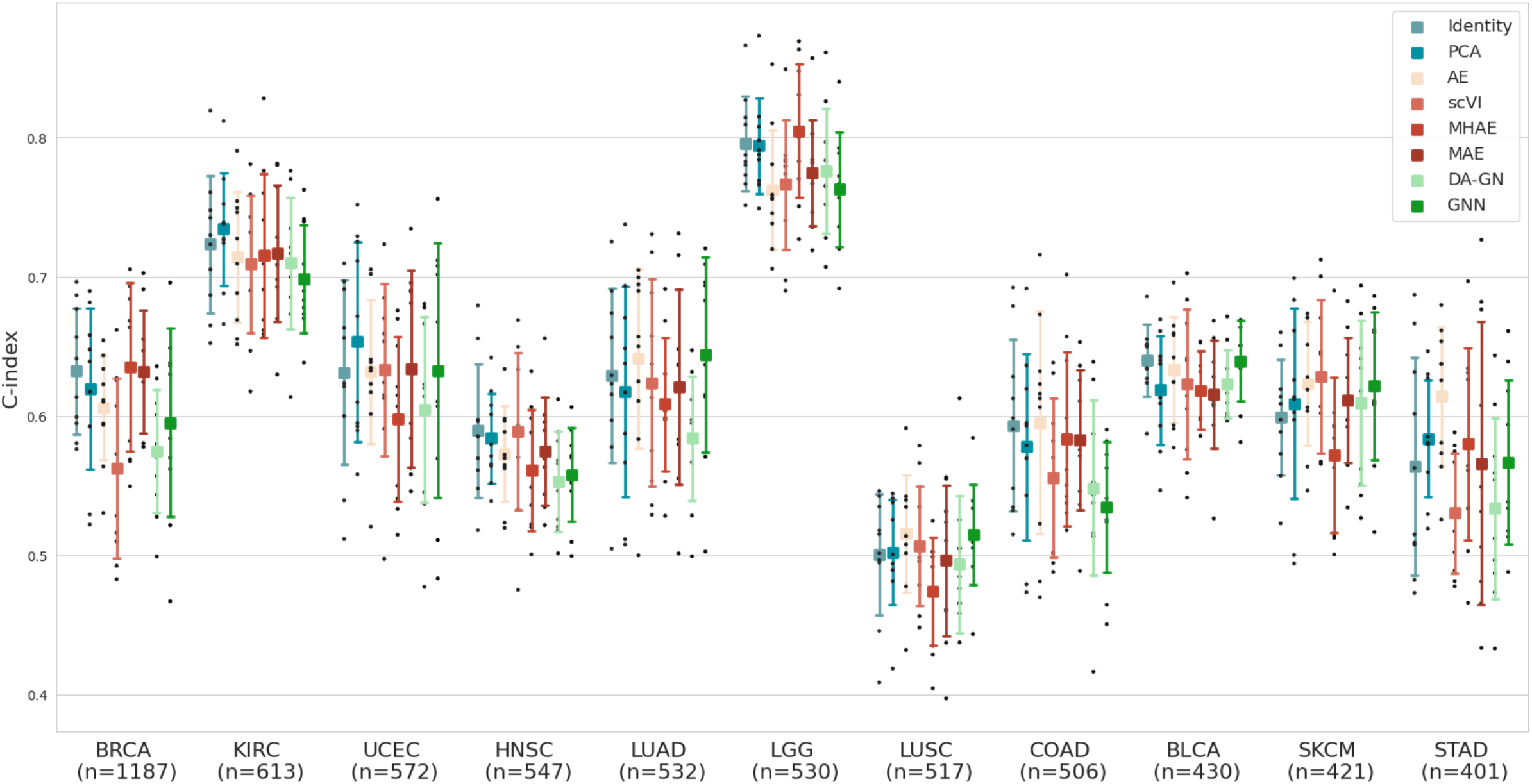
Comparison of performance on per-cohort OS prediction task on different TCGA cohorts for different bulk RNA-seq representation models. Black dots represent c-index results on different test folds. For each model, mean (square) c-index and standard deviation (intervals) are represented. Numbers on x axis labels represent the total number of samples available for this task in each cohort.

We also considered the OS prediction task in a pan-cancer setup (detailed in the Material and Methods section). For this task, most of our models, including baselines, produced comparable performances when taking into account test set variability. Even though results are not directly comparable because of different evaluation frameworks, State-of-the-art (SOTA) models such as MultiSurv (Vale-Silva et Rohr 2021) when using solely RNA-seq data exhibits comparable performance, 0.758 (0.735 - 0.780), to most of our models, including baselines. As shown in **Fig 2**, we observed that deep learning methods did not exhibit a clear advantage over all baselines, also in this pan-cancer case. Indeed, PCA, masked auto-encoders (MAE) and multi-head auto-encoders (MH-AE) equally emerged as the best-performing methods on pairwise comparison under the 75% criterion acceptance (**Supp. Figure S6**). Once again, mean c-indexes for all representation models differ for no more than 1.5%. Notice that in this case, similar conclusions could have been obtained by comparing mean c-index over test folds (cfr. Supp. **Table S5**).

**Figure 2.**
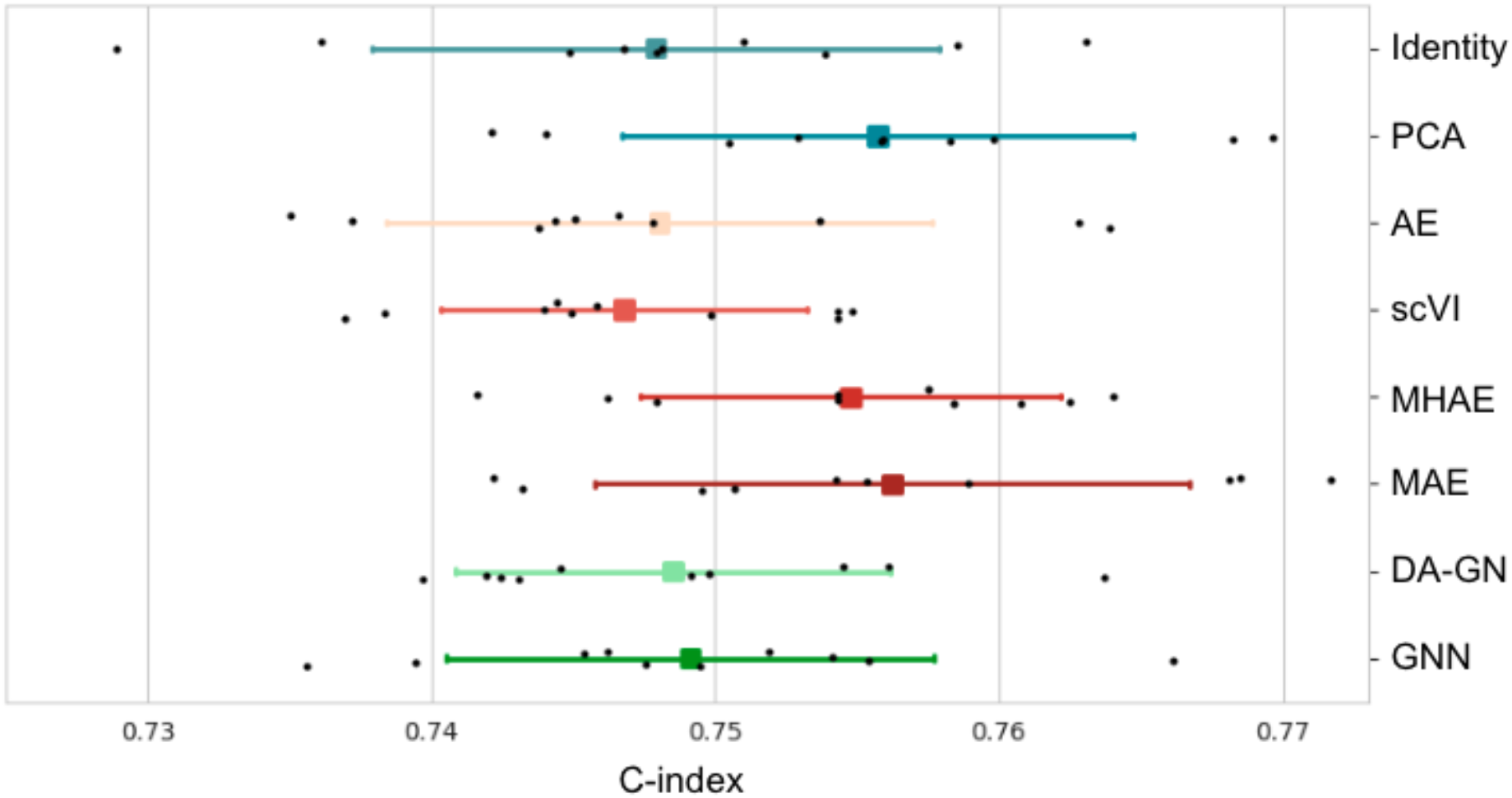
Comparison of performance on pan-cohort OS prediction task on TCGA dataset for different bulk RNA-seq representation models. Black dots represent c-index results on different test folds. For each model, mean (square) c-index and standard deviation (intervals) are represented.

The third task we considered was the Gene Essentiality prediction task. As expected, models obtained higher scores on the Overall Spearman correlation than the per-gene one (**Fig 3**), with differences between the best and the worst models around 0.6% and 8% for the overall and per-gene correlations respectively. Indeed, it is a more challenging task to grasp the distinction within a specific gene across various cell lines than it is to do so across all genes and cell lines. Interestingly, we see in this task that DRL methods more clearly demonstrated a superior performance compared to baselines on pairwise comparison under the 75% criterion acceptance (**Supp. Figure S7**). In particular, MAE and DA-GN proved to be the most effective approaches for both overall and per-gene correlations. Nonetheless, PCA reaches comparable results with best performing DRL methods, with differences of the order of 0.2% for overall correlation and 4% for per-gene correlation. Also in this case, mean Spearman correlation over test folds confirm superiority of both MAE and DA-GN over the other methods, and in general of DRL over simple baselines. While our evaluation framework and data are different from the original DeepDEP paper, preventing any direct comparison, we computed per-gene Pearson correlation scores to verify the coherence of our predictions to SOTA models’ performance ranges. We observed mean correlations per model ranging from 0.268, to 0.294, to compare with the 0.14 per-gene Pearson correlation score for Exp-DeepDEP (expression-only model) and 0.17 for DeepDep (on the 5 modalities).

**Figure 3.**
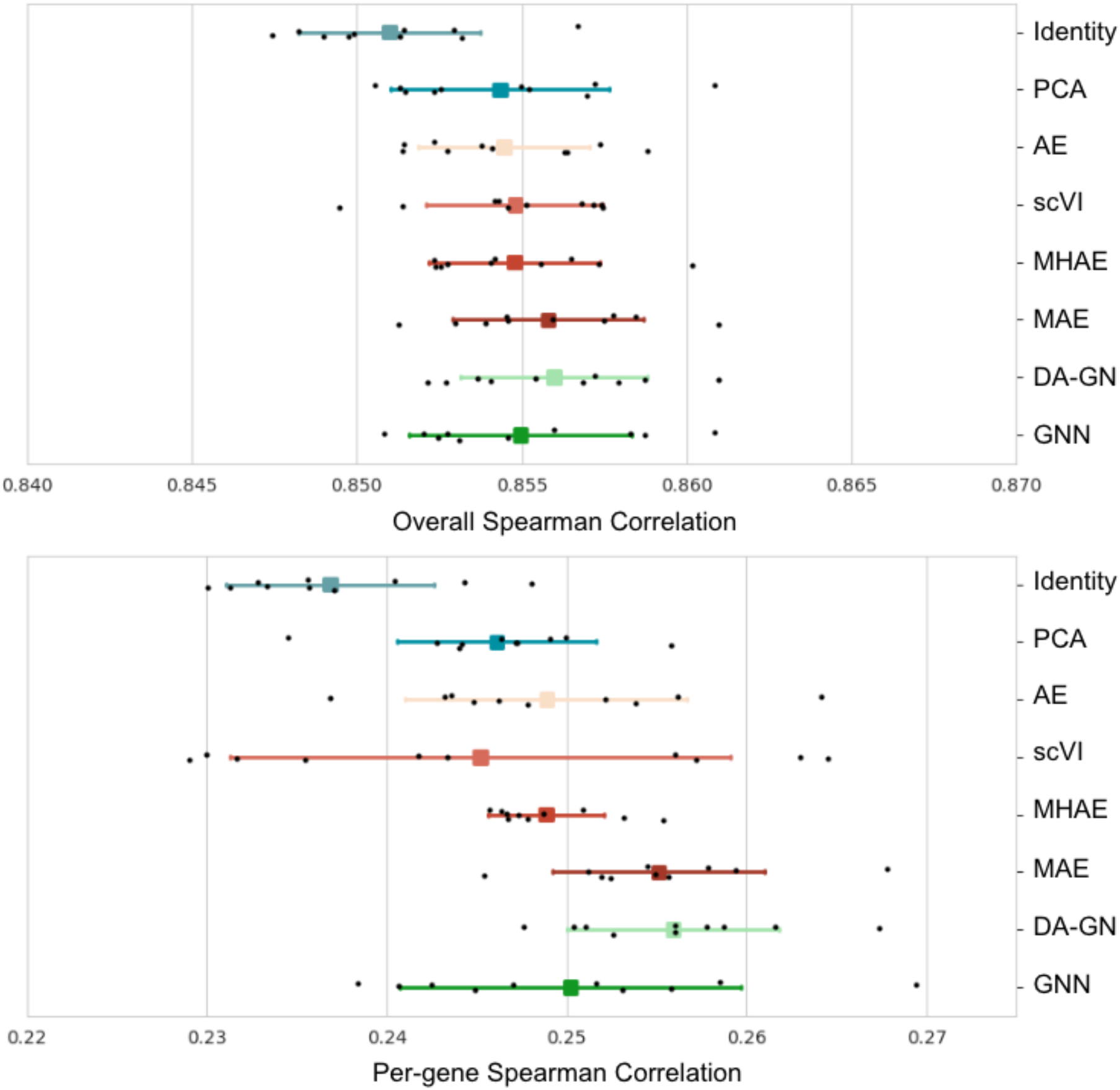
Comparison of performance on gene essentiality prediction task on DepMap dataset for different bulk RNA-seq representation models. Top panel) Spearman Rank correlation distributions between predicted and observed gene essentialities: black dots represent results on different test folds. For each model, mean (square) Spearman Rank and standard deviation (intervals) are represented. Bottom panel) Same as Top panel, but correlation computed per-gene.

All in all, we observed that no model clearly outperforms all the others for all tasks and all cohorts. Our study highlighted high variability of performances across different test sets for all tasks and models, showing the importance of adapting a rigorous evaluation framework taking these into account. Notably, we show that performing bootstrapping on a single test set could lead to false claims of superiority for certain models that are not generalizable to other data splits (**Fig 4a**).

**Figure 4.**
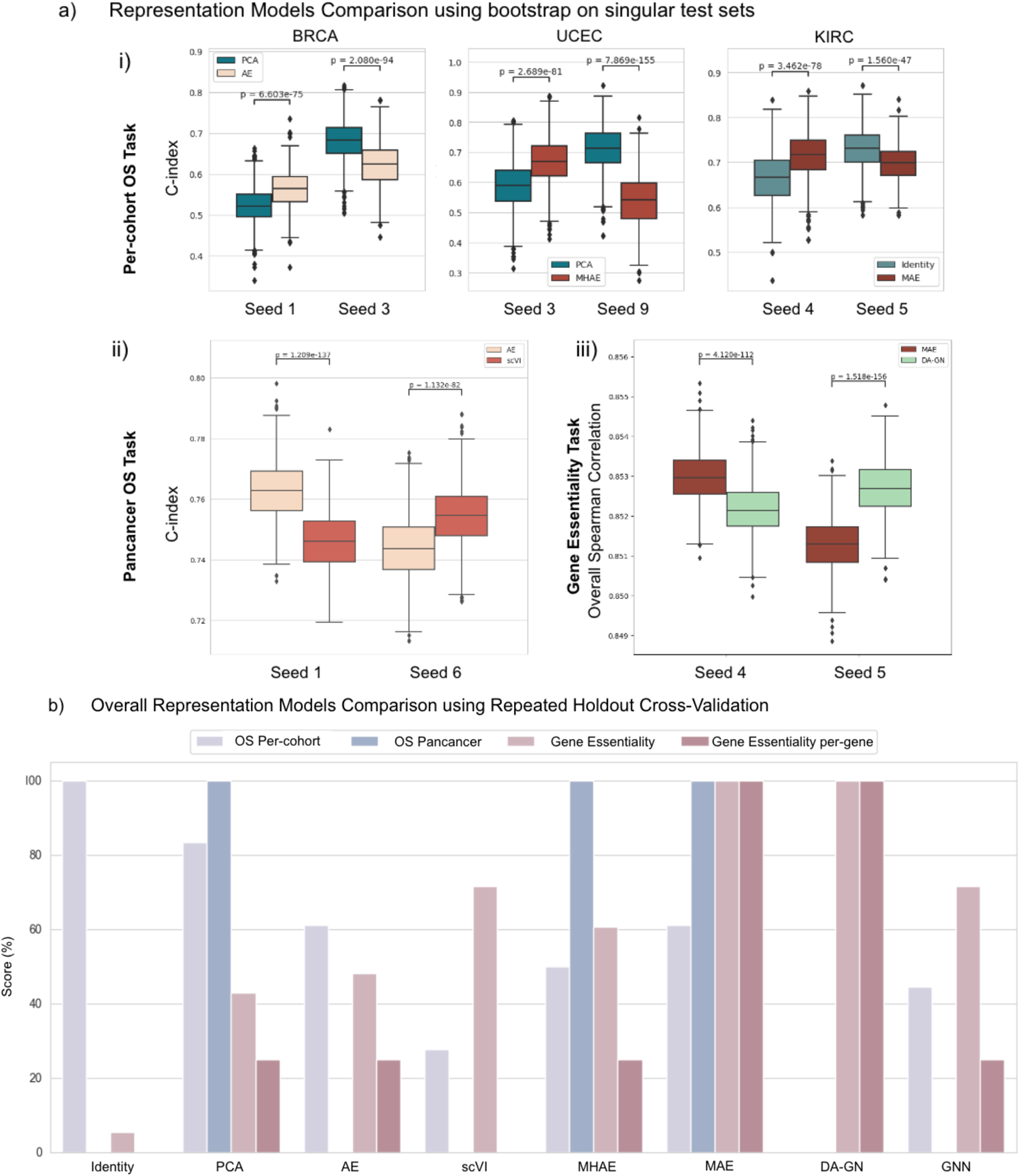
Comparison of model evaluation processes. **a) Test metric distributions obtained with bootstrap (n=1,000 per seed) on the test sets generated by different seeds on the downstream tasks.** Models were trained following our repeated holdout pipeline, but graphs show results only on arbitrary chosen individual test sets to showcase the limitations of this evaluation process. *p-values are computed using Wilcoxon tests.* i*) Comparison of representation models on 3 example cohorts from the per-cohort OS prediction task. ii) Comparison of AE and scVI on the pan-cancer task. iii) Comparison of MAE and DA-GN on the Gene Essentiality task.* **b) Overall comparison of the different representations models over the tasks Per-cohort OS Prediction, Pan-cohort OS Prediction and Gene Essentiality (Overall Metric and per-gene)**. *The score for a given model corresponds to the number of winning pairwise comparisons to other models according to our acceptance test criteria. The sum is then normalized by task to obtain the final score and is expressed as a percentage*.

Nonetheless, as described in the Methods section, the acceptance criterion can be used to define another scoring system to evaluate the relative superiority of a model over the others, that by construction is more and more consistent once increasing the number of test folds. The overall results about relative scoring of the models on the different tasks, using acceptance criterion and pairwise comparison, are shown in **Fig 4b**. Under this scoring system, the baseline representation models appear as the best models for survival prediction, Identity for Per-cohort task and PCA for Pan-cancer (together with MHAE and MAE). On the other hand, all DRL methods can be considered better (or at least equivalent) choices for Gene Essentiality prediction tasks, under both overall and per-gene correlation metrics.

### Pre-training influence on auto-encoders

Adding another dimension to our benchmark, we investigated the hypothesis that deep learning methods performances would benefit from pre-training on external datasets. For this use case, we focused on the auto-encoder based model on the per-cohort survival prediction and the gene essentiality prediction tasks. For both tasks, the external dataset for pre-training is taken from TCGA, removing the downstream task cohorts for the survival prediction. Specifically for the gene essentiality task, this setting allows us to potentially extend to RNA-seq-only models the claim from DeepDEP that a dataset representing different entities such as patient tumor samples can still be used to train a model to generate consistent embeddings of cell lines expression profiles.

We first repeated the survival prediction tasks in per-cohort TCGA datasets, by exploring the two different pretraining strategies described in the Methods section, on basic auto-encoders. As shown in **Fig 5** and **Supp Fig S8**, while in terms of mean performances, PreAE finetuned models outperform PreAE on almost all cohorts (with the exception of LUSC and SKMC), with differences of performances between AE and pretrained models ranging from 1 to 11% at most (**Table S3**), the superiority of PreAE finetuned is less clear in terms of 75% acceptance criterion on test folds.

**Figure 5.**
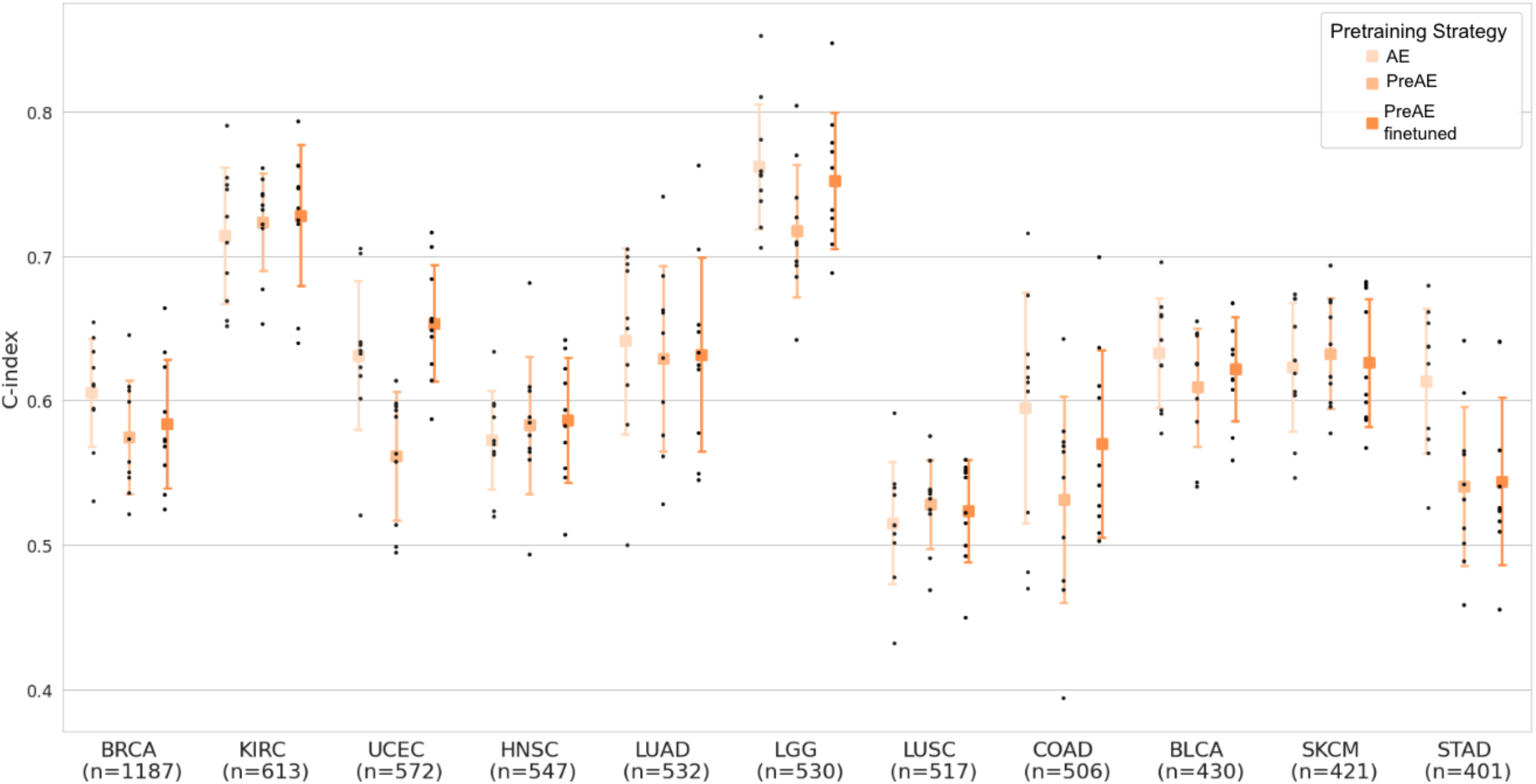
Impact of the pre-training strategy on the performance of autoencoders on the per-cohort OS prediction task on different TCGA bulk RNA-seq cohorts. Black dots represent c-index results on different test folds. Numbers on x axis labels represent the total number of samples available for this task in each cohort.

When considering the Gene Essentiality prediction task on the DepMap dataset, the results shown in **Fig 6** and **Supp. Fig S9** highlight that autoencoders pre-trained on TCGA samples (details in Methods section) struggle to outperform non pre-trained auto-encoders under the acceptance criterion assumed in this paper. This is the case for both pre-training strategies considered in this paper (with or without fine-tuning), and for both overall and per-gene correlations. Notice that PreAE (for which only the prediction model on top of the representation obtained on TCGA samples is trained on CCLE, the downstream task dataset) reaches performances which differ from the AE by 0.25% and 5% for the overall and per-gene correlation respectively. Moreover, retraining on CCLE dataset consistently improves the performance of PreAE for this task. It is to be noted that our setting here is different from DeepDEP as we do not focus on developing an end-to-end model integrating multiple modalities but we conducted similar experiments on Exp-DeepDEP using the original code baseline (see Supplementary Material for details) and reached similar conclusions (**Supp. Fig S10**).

**Figure 6.**
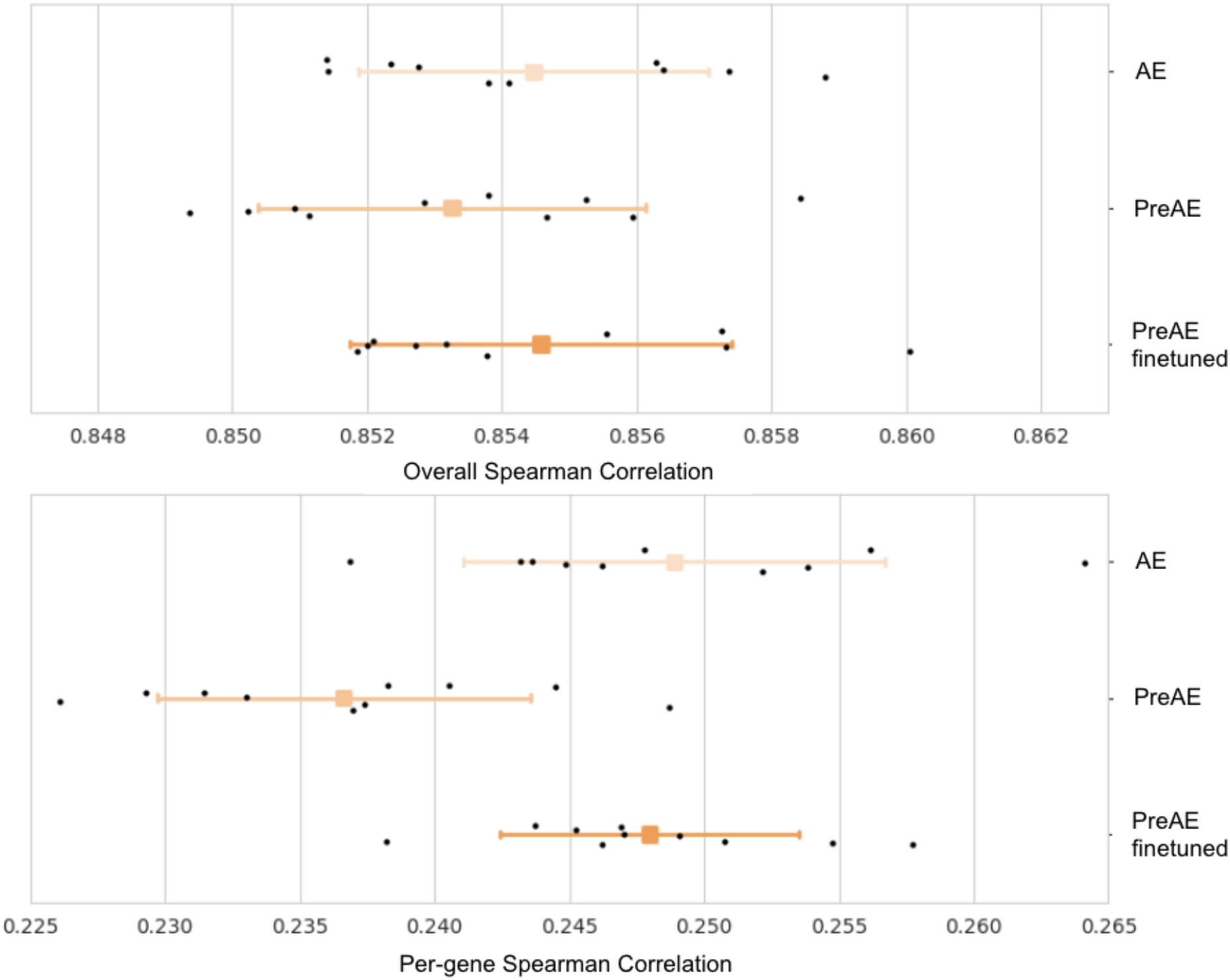
Impact of the pre-training strategy on TCGA samples on the performance of autoencoders on Gene Essentiality prediction task on DepMap dataset. *Top panel) Correlation distributions between predicted and observed gene essentialities: black dots represent results on different test folds. Bottom panel) Same as Top Left panel, but correlation computed per-gene*.

## Discussion

In this paper, we benchmarked different architectures for representation of bulk RNA-Seq data against two classical downstream tasks in cancer research: survival and gene essentiality prediction. Notably, we observed that simple methods for representation like Identity and PCA can achieve comparable or even superior performances with respect to more complex or deep models. In particular, we observed that simple baseline models can be considered as achieving superior performance over other DRL on survival prediction tasks on patient TCGA data, while DRL methods seem to have a slight but consistent advantage over baselines on gene essentiality prediction tasks on cell line DepMap data. Moreover, our findings indicated consistent improvements across tasks in auto-encoders (AE) when incorporating masking and multi-head techniques, suggesting promising avenues for future research.

Furthermore, we utilized our benchmarking framework to investigate the impact of pre-training on the same models and tasks. We found that pre-trained autoencoders (AE) on patient samples (TCGA) generated representations that achieved comparable gene essentiality performance on cell-lines data to fine-tuned methods, without any specific preprocessing or alignment between TCGA and CCLE datasets. This shows that the representation is able to transfer knowledge and suggests potential for significant computational time savings by pretraining a “feature extractor”, a practice common in image processing. However, we observed that pre-training did not lead to significant performance improvements in both overall survival (OS) and gene essentiality (GE) prediction, even after subsequent fine-tuning.

We also showed that comparing models on a singular test set sampled from the task datasets can lead to misinterpretations even if coupled with statistical significance testing. This highlights the importance of adopting standardized evaluation practices to avoid unwarranted claims of superiority for novel techniques. This seems particularly true when training representation models on patient bulk RNA-seq data, and is probably due to the poor ratio samples/features and the intrinsic complexity of the underlying biology.

In this study, we tackled the issue of reproducibility of results in the field of deep learning on omics data for cancer research. We focused on building a robust, reproducible and fair benchmarking framework to compare different machine learning models built on bulk RNA-seq data with respect to their performance on different downstream tasks. The key elements used in the pipeline we developed (repeated holdout cross-validation and fixed HP tuning budget) are not new per-se (Filiot et al. 2023) but to our knowledge this framework has been used in a limited number of studies (Smith et al. 2020) evaluating ML models on omics data in cancer-specific downstream tasks. Tuning directly on a cross-validation without a held out test set could lead to inflated performance and justifies the need for a nested cross-validation framework or the proposed repeated holdout test set. We chose the repeated holdout test to be able to provide confidence intervals and to provide a fairer metric (see details in Supplementary Materials). Additionally, we adopted the acceptance criterion from (Varoquaux et Colliot 2023) to compare model performance, in order to focus on meaningful improvements on finite-sized test sets. This criterion allows us to establish a scoring system for relative model performance, complementing standard mean performance considerations by accounting for variations from non-model factors like random splits during cross-validation and HP tuning strategy. We applied this benchmarking framework to evaluate the performance of different dimensionality reduction methods, including various model architectures and training techniques, on different downstream tasks (survival on patient data and gene essentiality on cell lines) using TCGA and DepMap public pan-cancer datasets.

We focused this work on fair comparisons and ablation studies, ensuring that each model was tested independently without combining different elements such as scaling, normalization, and gene selection. This approach allowed us to identify the strengths and weaknesses of each model more accurately. We believe this framework to be robust and generalizable to other tasks. For instance, while in the original VIME paper (Yoon et al. 2020) it was observed that the addition of mask prediction as a subtask provided improved performances, in our specific use case our ablation study revealed that once applied on bulk RNA-seq data, this addition actually worsened the models’ performance and increased training time.

While our study provides valuable insights, we acknowledge some limitations. For instance, we chose not to incorporate multi-omics data in our analysis due to prior research (Vale-Silva et Rohr 2021; Chiu et al. 2021) indicating marginal gains or lack of statistical significance when integrating such data in related studies. The framework uses a fixed budget HP tuning, which could arguably have been different per model. Larger models and higher HP budgets could potentially yield better performances. Our study also focused on breadth rather than depth, exploring many models without deep parameter studies and tuning. Regarding the comparison with a related method, DeepDEP, we observed that the datasets used in our study were not identical, possibly leading to adaptations in hyperparameters. Nonetheless, we demonstrated that pre-training did not consistently improve performance, supporting our earlier observations. Lastly, we want to highlight that the highest-performing method on a single metric may not necessarily be the most effective approach in practical applications. Taking into account other relevant considerations, such as model interpretability, can contribute to more informed decisions regarding method selection for specific use cases.

Future development could include extending our tasks by considering out-of-domain evaluation metrics or exploring additional tasks like drug response prediction. The computational scale of the study can be increased, with greater budget for HP tuning and scaling up the number of test sets, and combining models for potentially greater improvements is a natural step. Additionally, different architectures from the regular auto-encoders could potentially leverage the pretraining process more efficiently as well as pretraining on larger bulk RNA-seq datasets (Lachmann et al. 2018). Recent learnings from large pretrained models applied to single-cell data could be applied to our use case, such as different feature preprocessing of RNA-seq (Theodoris et al. 2023; Cui et al. 2023).

In conclusion, we believe our study brings valuable insights to the understanding of deep learning methods for bulk RNA-seq data representations in cancer research. By providing a robust evaluation framework we aim to facilitate fair comparisons of novel approaches against established methods, thus advancing the field towards more reliable and impactful techniques. The limitations identified in this study also offer guidance for future research directions and potential refinements to enhance the application of deep learning in cancer research.

## Data and Code Availability

The cancer TCGA data was downloaded from recount3 https://rna.recount.bio/, the associated clinical data from TCGA-CDR^1^ and the cell lines datasets from the DepMap portal https://depmap.org/portal/. The code is available in a public repository provided upon publication.

## Contributions

B.G., An.D., V.C, V.K., M.B., Y.D and A.R designed the evaluation framework and the different tasks; An.D, B.G, V.K, G.D, V.C, J.E.K, K.O, S.G., Al.D, L.H., R.G., R.L and C.E wrote the code to develop the evaluation pipeline; B.G, An.D, V.C, V.K, J.E.K, G.D, S.G, K.O., Al.D, R.L, C.E, Y.D, A.R analyzed the results; Y.D., A.R. and E.D. coordinated and supervised the work; B.G., An.D., V.C, Y.D, E.D. and A.R wrote the paper with the assistance and feedback of all the other co-authors. All authors reviewed and approved the final manuscript.

## Acknowledgements

The results shown here are in whole or part based upon data generated by the TCGA Research Network: https://www.cancer.gov/tcga. We thank Oussama Tchita and Omar Darwiche Domingues for their valuable contributions and comments to strengthen our pipeline and coding best practices. We thank Gilles Wainrib for initial ideas and discussions, Nicolas Loiseau for his advice and statistical expertise, Floriane Montanari, Benoît Schmauch, Gilles Wainrib and Jean-Philippe Vert for their detailed proofreading and insightful comments.

## Footnotes

1 https://gdc.cancer.gov/about-data/publications/PanCan-Clinical-2018

