## Supplementary Materials for "Robust Evaluation of Deep Learning-based Representation Methods for Survival and Gene Essentiality Prediction on Bulk RNA-seq Data"

#### Details on models hyperparameters

Table S1: Hyperparameters Ranges for Representation Models

<table>
<tr><th rowspan="2">Representation Model</th><th rowspan="2">Hyperparameter</th><th colspan="2">Ranges</th></tr>
<tr><th>OS Tasks</th><th>GE Task</th></tr>
<tr><td>PCA</td><td>Representation Dimension</td><td>⟦4, 256⟧</td><td>⟦16, 256⟧</td></tr>
<tr><td>AE</td><td>Representation Dimension</td><td>⟦4, 256⟧</td><td>⟦16, 256⟧</td></tr>
<tr><td></td><td>Hidden Units First Layer</td><td colspan="2">⟦256, 1024⟧</td></tr>
<tr><td></td><td>Additional Hidden Layers</td><td>⟦0, 1⟧</td><td>⟦0, 2⟧</td></tr>
<tr><td></td><td>Hidden Decrease Rate</td><td colspan="2">{0.5; 1}</td></tr>
<tr><td></td><td>Learning Rate</td><td>[5e-5, 5e-3]</td><td>[5e-6, 5e-4]</td></tr>
<tr><td></td><td>Batch Size</td><td>256</td><td>1024</td></tr>
<tr><td></td><td>Dropout Rate</td><td colspan="2">[0, 0.2]</td></tr>
<tr><td></td><td>Maximum Number of Epochs</td><td colspan="2">300</td></tr>
<tr><td></td><td>Early Stopping Patience</td><td colspan="2">50</td></tr>
<tr><td></td><td>Early Stopping Delta</td><td colspan="2">1e-3</td></tr>
<tr><td>scVI</td><td>Representation Dimension</td><td>⟦4, 256⟧</td><td>⟦16, 256⟧</td></tr>
<tr><td></td><td>Hidden Units First Layer</td><td colspan="2">⟦256, 1024⟧</td></tr>
<tr><td></td><td>Additional Hidden Layers</td><td>⟦0, 1⟧</td><td>⟦0, 2⟧</td></tr>
<tr><td></td><td>Hidden Decrease Rate</td><td colspan="2">{0.5; 1}</td></tr>
<tr><td></td><td>Learning Rate</td><td colspan="2">0</td></tr>
<tr><td></td><td>Batch Size</td><td>256</td><td>1024</td></tr>
<tr><td></td><td>Dropout Rate</td><td colspan="2">0.1</td></tr>
<tr><td></td><td>Maximum Number of Epochs</td><td colspan="2">Default scVI value</td></tr>
<tr><td>MAE</td><td>Representation Dimension</td><td colspan="2">256</td></tr>
<tr><td></td><td>Hidden Units First Layer</td><td colspan="2">512</td></tr>
<tr><td></td><td>Additional Hidden Layers</td><td colspan="2">0</td></tr>
<tr><td></td><td>Learning Rate</td><td>[5e-5, 5e-3]</td><td>[5e-5, 5e-4]</td></tr>
<tr><td></td><td>Batch Size</td><td>256</td><td>1024</td></tr>
<tr><td></td><td>Dropout Rate</td><td colspan="2">0</td></tr>
<tr><td></td><td>Corruption Probability</td><td colspan="2">[0.1, 0.5]</td></tr>
<tr><td></td><td>Maximum Number of Epochs</td><td colspan="2">1,000</td></tr>
<tr><td></td><td>Early Stopping Patience</td><td colspan="2">20</td></tr>
<tr><td></td><td>Early Stopping Delta</td><td colspan="2">1e-5</td></tr>
<tr><td>MHAE</td><td>Representation Dimension</td><td colspan="2">256</td></tr>
<tr><td></td><td>Hidden Units First Layer</td><td colspan="2">512</td></tr>
<tr><td></td><td>Additional Hidden Layers</td><td colspan="2">0</td></tr>
<tr><td></td><td>Learning Rate</td><td colspan="2">[5e-5, 5e-3]</td></tr>
<tr><td></td><td>Batch Size</td><td>256</td><td>1024</td></tr>
<tr><td></td><td>Dropout Rate</td><td colspan="2">{0; 0.1}</td></tr>
<tr><td></td><td>Maximum Number of Epochs</td><td colspan="2">1,000</td></tr>
<tr><td></td><td>Early Stopping Patience</td><td colspan="2">50</td></tr>
<tr><td></td><td>Early Stopping Delta</td><td colspan="2">1e-5</td></tr>
<tr><td></td><td>Auxiliary Head Neurons</td><td>Per-cohort: {32; 128}<br>Pancancer: 128</td><td>32</td></tr>
<tr><td></td><td>Auxiliary Head Dropout Rate</td><td>{0; 0.1}</td><td>0</td></tr>
<tr><td></td><td>Loss Weight for Aux. Head $\beta$</td><td>[0.001, 1]</td><td>[0.1, 10]</td></tr>
<tr><td>GNN</td><td>Representation Dimension</td><td colspan="2">128</td></tr>
<tr><td></td><td>Learning Rate</td><td colspan="2">1e-3</td></tr>
<tr><td></td><td>Batch Size</td><td colspan="2">32</td></tr>
<tr><td></td><td>Number of Epochs</td><td colspan="2">50</td></tr>
<tr><td></td><td>Message Passing</td><td colspan="2">$\{GraphConv; SAGE\}$</td></tr>
<tr><td></td><td>Channels</td><td colspan="2">{[8]; [8, 16]; [8, 16, 16]}</td></tr>
<tr><td></td><td>Louvain Cluster Resolution</td><td colspan="2">[10, 500]</td></tr>
<tr><td></td><td>Pooling</td><td colspan="2">$\{avg, max\}$</td></tr>
<tr><td></td><td>StringDB Threshold</td><td colspan="2">[0.7, 0.99]</td></tr>
<tr><td></td><td>Graph Encoder Dropout</td><td colspan="2">0.3</td></tr>
<tr><td></td><td>Decoder Hidden Layers</td><td colspan="2">{1, 2}</td></tr>
<tr><td></td><td>Decoder Hidden Decrease Rate</td><td colspan="2">0.5</td></tr>
<tr><td></td><td>Decoder Dropout Rate</td><td colspan="2">0.2</td></tr>
</table>

Table S1: Hyperparameters Ranges for Representation Models

<table>
<tr><th rowspan="2">Representation Model</th><th rowspan="2">Hyperparameter</th><th colspan="2">Ranges</th></tr>
<tr><th>OS Tasks</th><th>GE Task</th></tr>
<tr><td>DA-GN</td><td>Representation Dimension</td><td colspan="2">256</td></tr>
<tr><td></td><td>Hidden Units First Layer</td><td colspan="2">512</td></tr>
<tr><td></td><td>Additional Hidden Layers</td><td colspan="2">0</td></tr>
<tr><td></td><td>Learning Rate</td><td>[5e-5, 5e-3]</td><td>[5e-5, 5e-4]</td></tr>
<tr><td></td><td>Batch Size</td><td>256</td><td>1024</td></tr>
<tr><td></td><td>Dropout Rate</td><td colspan="2">0.1</td></tr>
<tr><td></td><td>Maximum Number of Epochs</td><td colspan="2">1,000</td></tr>
<tr><td></td><td>Early Stopping Patience</td><td colspan="2">20</td></tr>
<tr><td></td><td>Early Stopping Delta</td><td colspan="2">1e-5</td></tr>
<tr><td></td><td>Noise Level</td><td colspan="2">[0.01, 1]</td></tr>
<tr><td></td><td>Number of copies</td><td colspan="2">4</td></tr>
<tr><td>PreAE</td><td>Representation Dimension</td><td colspan="2">⟦16, 256⟧</td></tr>
<tr><td></td><td>Hidden Units First Layer</td><td colspan="2">⟦256, 1024⟧</td></tr>
<tr><td></td><td>Additional Hidden Layers</td><td colspan="2">⟦0, 2⟧</td></tr>
<tr><td></td><td>Hidden Decrease Rate</td><td colspan="2">{0.5; 1}</td></tr>
<tr><td></td><td>Learning Rate</td><td colspan="2">[5e-6, 5e-4]</td></tr>
<tr><td></td><td>Batch Size</td><td colspan="2">1024</td></tr>
<tr><td></td><td>Dropout Rate</td><td colspan="2">[0, 0.2]</td></tr>
<tr><td></td><td>Maximum Number of Epochs</td><td colspan="2">1,000</td></tr>
<tr><td></td><td>Early Stopping Patience</td><td colspan="2">20</td></tr>
<tr><td></td><td>Early Stopping Delta</td><td colspan="2">1e-5</td></tr>
</table>

**Table S1** Hyperparameters Range for Representation Models. Brackets represent sets of values, single [ ] represent float intervals and double [[ ]] represent integer ranges

### Details on representation models implementation

We include here more details on the implementation of different representations models and meaning of certain hyperparameters names. Models not mentioned below are considered described thoroughly in the main text.

#### Auto-Encoders

In our implementation, Hidden Units First Layers correspond to the number of neurons in the first layer after the input in the AE-based architectures (AE, PreAE, scVI, MAE, MHAE, DA-GN). The Additional Hidden Layers correspond to the number of layers in the Encoder / Decoder excluding the representation layer. The Hidden Decrease Rate controls the bottleneck of the Encoder : a value of 0.5 means that at each additional hidden layer, the number of neurons is divided by 2. The batch size was fixed and not used as a hyperparameter following advice from (Godbole et al. 2023).

#### Masking Auto-Encoders

In VIME, the authors introduce an innovative masking scheme (compared to Gaussian noise addition or binary masking), in which they :

1. Generate a permuted variant of the samples

2. Generate a binary mask
3. Compose the binary mask and the permutation to generate a corrupted sample $x_tilde = m * x_perm + (1-m) * x$, where $x$ is the original samples, $m$ a binary mask sampled from a Bernouilli distribution and $x_perm$ the permuted sample.

#### Multi-Head Auto-Encoders

The MH auto-encoder simplified architecture is depicted below:

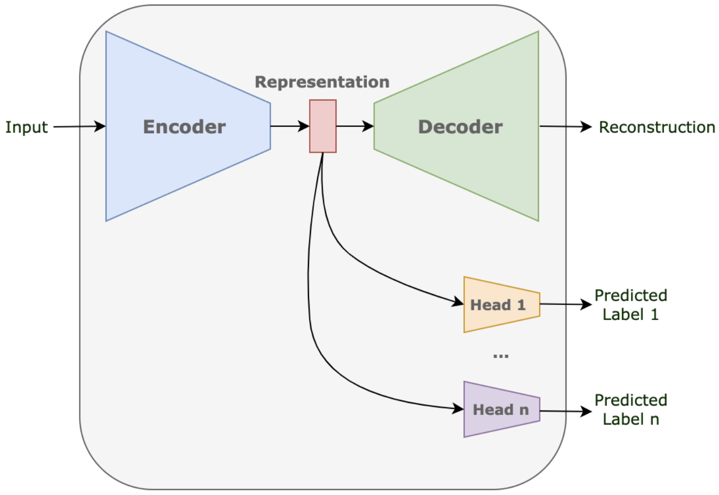


This model was trained using a two-term loss function:

$$\mathcal{L} = \mathcal{L}_{rec} + \beta * \mathcal{L}_{aux},$$

where $\beta$ is the hyperparameter controlling the weight between the two terms, $\mathcal{L}_{rec}$ is the auto-encoder reconstruction error (mean squared error), and $\mathcal{L}_{aux}$ is the auxiliary head loss function. The latter depends on the predicted endpoint: mean squared error for gene essentiality, and cox loss for overall survival.

#### Graph Neural Networks

The STRING (Szklarczyk et al. 2023) data was preprocessed by keeping only genes present in our omics data and forming the induced subgraph based on this gene list. Additionally, we retained only the most confident interactions, using the 'combined score' column, by setting a quantile parameter q. The quantile value ranged from 0.7 (as suggested by the STRING db) to 0.99. Nodes not belonging to the largest connected component were discarded to ensure downstream clustering did not include isolated single nodes.

Clustering was then performed on the graph, which would be used in the pooling part of the encoder. Our goal was to define tightly connected gene communities within the graph. The Louvain algorithm implemented in networkX was employed to detect these communities, with the resolution parameter controlling the granularity of the clusters. These clusters, presented as gene lists, were given as input to our GNN encoder, along with the actual graph and the RNA-seq data.

Our GNN model was created using the Pytorch Geometric library. It was built as an auto-encoder, comprising a GNN encoder and a classical MLP decoder. The model aimed to reconstruct the omics signal given the omics signal and the graph as input. The encoder consisted of stacked classical GNN layers (SAGEconv (Hamilton, Ying, et Leskovec 2018), GraphConv(Morris et al. 2021)), typically ranging from 1 to 3 layers. The number of channels (i.e., size of the node embedding) increased at each layer, with different values tested, such as [8,8,16], to obtain a deep node embedding of dimension $d$. To reduce the dimensionality of the embeddings, nodes were grouped per cluster defined in the previous paragraph. Cluster-level representations were generated by applying pooling layers (average pooling or max pooling) to genes within the same cluster. This resulted in $n_cluster$ $d$-dimensional embeddings, which were concatenated into a single $n_cluster x d$ dimensional embedding. To ensure comparability with other models and determine the embedding dimension, this embedding was passed through a single MLP layer.

The decoder, derived from our Auto-Encoder architecture, was concatenated with the encoder. The entire model was trained end-to-end to reconstruct the bulk RNA-seq signal when provided with the expression data and graph inputs.

#### Exp-DeepDEP Experiments

In our setting for the Gene Essentiality task, we focused mostly on the cell lines RNA-seq representations but did not fine-tune the fingerprints representation nor created an end-to-end deep learning model specifically for this task such as DeepDEP. DeepDEP is a multimodal model that takes into account not only bulkRNA-seq data but also mutations, copy number alterations, methylation data and fingerprints, integrating them through the combination of different encoder heads in the architecture of the model. They show that, with pretraining on TCGA, this model improves performance in their evaluation framework compared to no pretraining. They also show that a simplified version of their model, Exp-DeepDEP, obtained similar performances as DeepDEP using only RNA-seq and fingerprints. With the pre-trained auto-encoders presented above, we investigated whether a pre-trained representation model on TCGA could provide better embeddings for the expression profiles of the downstream task dataset when concatenated with fixed fingerprints representations and passed through the prediction model (LGBM) directly. Similarly to their work, we selected the top 5,000 variable genes on TCGA and trained an end-to-end Exp-DeepDEP on our data splits, with and without the pretraining step, to investigate the influence of pre-training in the same setting as the original paper when focused only on expression data. The pretraining step was performed similarly as the original Exp-DeepDEP and the same hyperparameters. After pretraining, Ex-DeepDEP models were trained on CCLE following our data splits using again the same HPs as the original paper. To do so, we downloaded the original code from DeepDEP on CodeOcean[2] , modified it and made it available in our GitHub repository.

[2] https://codeocean.com/capsule/7914207/tree/

### Details on prediction models hyperparameters

Table S2: Hyperparameters Ranges for Prediction Models

| Prediction Model | Hyperparameter | Range |
|---|---|---|
| MLP (OS Tasks) | Hidden Layers | [128] |
| | Learning Rate | [1e-5, 1e-3] |
| | Batch Size | 256 |
| | Dropout Rate | 0 |
| | Maximum Number of Epochs | 1,000 |
| | Early Stopping Patience | 50 |
| | Early Stopping Delta | 1e-5 |
| LGBM (Gene Essentiality) | Learning Rate | [0.01, 0.3] |
| | L1 Regularization | [0, 100] |

**Table S2** Hyperparameters Range for Prediction Models. Brackets represent sets of values, single [ ] represent float intervals and double [[ ]] represent integer ranges

#### Results Tables

**Table S3** Test sets c-index statistics for the per-cohort OS prediction task. Best performance based on mean results are shown in boldface.

| Cohort | Representation Model | Mean Metric | Median Metric | Standard Deviation |
|---|---|---|---|---|
| BRCA | Identity | 0,632 | 0,627 | 0,045 |
| | PCA | 0,619 | 0,628 | 0,058 |
| | AE | 0,606 | 0,611 | 0,037 |
| | scVI | 0,562 | 0,550 | 0,065 |
| | MHAE | **0,635** | 0,666 | 0,060 |
| | MAE | 0,632 | 0,631 | 0,044 |
| | DA-GN | 0,575 | 0,575 | 0,044 |
| | GNN | 0,595 | 0,607 | 0,068 |
| KIRC | Identity | 0,723 | 0,727 | 0,049 |
| | PCA | **0,734** | 0,733 | 0,040 |
| | AE | 0,714 | 0,718 | 0,047 |
| | scVI | 0,709 | 0,708 | 0,049 |
| | MHAE | 0,715 | 0,700 | 0,059 |
| | MAE | 0,717 | 0,712 | 0,049 |
| | DA-GN | 0,710 | 0,712 | 0,047 |
| | GNN | 0,698 | 0,686 | 0,039 |
| UCEC | Identity | 0,631 | 0,640 | 0,066 |
| | PCA | **0,653** | 0,641 | 0,072 |
| | AE | 0,631 | 0,633 | 0,052 |
| | scVI | 0,633 | 0,639 | 0,062 |
| | MHAE | 0,598 | 0,611 | 0,059 |
| | MAE | 0,633 | 0,638 | 0,071 |
| | DA-GN | 0,604 | 0,617 | 0,067 |
| | GNN | 0,632 | 0,639 | 0,091 |
| HNSC | Identity | **0,589** | 0,585 | 0,048 |
| | PCA | 0,584 | 0,575 | 0,032 |
| | AE | 0,573 | 0,571 | 0,034 |
| | scVI | **0,589** | 0,590 | 0,056 |
| | MHAE | 0,561 | 0,554 | 0,043 |
| | MAE | 0,575 | 0,571 | 0,039 |
| | DA-GN | 0,553 | 0,556 | 0,036 |
| | GNN | 0,558 | 0,563 | 0,033 |
| LUAD | Identity | 0,629 | 0,626 | 0,062 |

| | | | | |
|---|---|---|---|---|
| | PCA | 0,618 | 0,627 | 0,075 |
| | AE | 0,641 | 0,653 | 0,065 |
| | scVI | 0,624 | 0,625 | 0,074 |
| | MHAE | 0,608 | 0,605 | 0,048 |
| | MAE | 0,621 | 0,630 | 0,070 |
| | DA-GN | 0,584 | 0,588 | 0,045 |
| | GNN | **0,644** | 0,660 | 0,070 |
| LGG | Identity | 0,796 | 0,785 | 0,034 |
| | PCA | 0,794 | 0,789 | 0,034 |
| | AE | 0,762 | 0,756 | 0,043 |
| | scVI | 0,766 | 0,779 | 0,047 |
| | MHAE | **0,805** | 0,803 | 0,048 |
| | MAE | 0,775 | 0,777 | 0,038 |
| | DA-GN | 0,776 | 0,771 | 0,045 |
| | GNN | 0,763 | 0,766 | 0,041 |
| LUSC | Identity | 0,500 | 0,508 | 0,044 |
| | PCA | 0,502 | 0,497 | 0,038 |
| | AE | 0,515 | 0,514 | 0,042 |
| | scVI | 0,507 | 0,501 | 0,043 |
| | MHAE | 0,474 | 0,484 | 0,039 |
| | MAE | 0,496 | 0,517 | 0,054 |
| | DA-GN | 0,493 | 0,480 | 0,049 |
| | GNN | 0,515 | 0,513 | 0,036 |
| COAD | Identity | 0,593 | 0,594 | 0,062 |
| | PCA | 0,578 | 0,586 | 0,067 |
| | AE | **0,595** | 0,614 | 0,080 |
| | scVI | 0,556 | 0,562 | 0,057 |
| | MHAE | 0,584 | 0,556 | 0,063 |
| | MAE | 0,583 | 0,580 | 0,050 |
| | DA-GN | 0,548 | 0,548 | 0,063 |
| | GNN | 0,534 | 0,536 | 0,047 |
| BLCA | Identity | **0,640** | 0,642 | 0,026 |
| | PCA | 0,619 | 0,625 | 0,039 |
| | AE | 0,633 | 0,633 | 0,038 |
| | scVI | 0,623 | 0,627 | 0,054 |
| | MHAE | 0,618 | 0,615 | 0,028 |
| | MAE | 0,615 | 0,614 | 0,039 |

| | | | | |
|---|---|---|---|---|
| | DA-GN | 0,623 | 0,619 | 0,024 |
| | GNN | 0,639 | 0,645 | 0,029 |
| SKCM | Identity | 0,599 | 0,603 | 0,041 |
| | PCA | 0,609 | 0,613 | 0,068 |
| | AE | 0,623 | 0,623 | 0,045 |
| | scVI | 0,628 | 0,626 | 0,055 |
| | MHAE | 0,572 | 0,576 | 0,056 |
| | MAE | 0,611 | 0,604 | 0,045 |
| | DA-GN | 0,609 | 0,626 | 0,059 |
| | GNN | 0,621 | 0,615 | 0,053 |
| STAD | Identity | 0,564 | 0,539 | 0,078 |
| | PCA | 0,584 | 0,580 | 0,042 |
| | AE | **0,614** | 0,631 | 0,050 |
| | scVI | 0,530 | 0,522 | 0,043 |
| | MHAE | 0,580 | 0,573 | 0,069 |
| | MAE | 0,566 | 0,540 | 0,102 |
| | DA-GN | 0,534 | 0,533 | 0,065 |
| | GNN | 0,567 | 0,552 | 0,059 |

**Table S4** Test sets c-index statistics for the pan-cancer OS prediction task. Best performance based on mean results are shown in boldface.

| Representation Model | Mean Metric | Median Metric | Standard Deviation |
|---|---|---|---|
| Identity | 0,748 | 0,748 | 0,010 |
| PCA | **0,756** | 0,756 | 0,009 |
| AE | 0,748 | 0,746 | 0,010 |
| scVI | 0,747 | 0,745 | 0,006 |
| MHAE | 0,755 | 0,756 | 0,007 |
| MAE | **0,756** | 0,755 | 0,010 |
| DA-GN | 0,748 | 0,747 | 0,008 |
| GNN | 0,749 | 0,748 | 0,009 |

**Table S5** Test sets overall spearman correlation statistics for the gene essentiality task. Best performance based on mean results are shown in boldface.

| Representation Model | Mean Metric | Median Metric | Standard Deviation |
|---|---|---|---|
| Identity | 0,8510 | 0,8506 | 0,0028 |
| PCA | 0,8543 | 0,8538 | 0,0033 |
| AE | 0,8545 | 0,8539 | 0,0026 |
| scVI | 0,8548 | 0,8549 | 0,0027 |
| MHAE | 0,8548 | 0,8541 | 0,0026 |
| MAE | 0,8558 | 0,8553 | 0,0029 |
| DA-GN | **0,8560** | 0,8561 | 0,0028 |
| GNN | 0,8550 | 0,8538 | 0,0034 |

**Table S6** Test sets per-gene spearman correlation statistics for the gene essentiality task. Best performance based on mean results are shown in boldface.

| Representation Model | Mean Metric | Median Metric | Standard Deviation |
|---|---|---|---|
| Identity | 0,237 | 0,236 | 0,006 |
| PCA | 0,246 | 0,247 | 0,006 |
| AE | 0,249 | 0,247 | 0,008 |
| scVI | 0,245 | 0,243 | 0,014 |
| MHAE | 0,249 | 0,248 | 0,003 |
| MAE | 0,255 | 0,255 | 0,006 |
| DA-GN | **0,256** | 0,256 | 0,006 |
| GNN | 0,250 | 0,249 | 0,009 |

**Table S7** Test sets c-index statistics for the per-cohort OS prediction task for pretrained experiments. Best performance based on mean results are shown in boldface.

| Cohort | Representation Model | Mean Metric | Median Metric | Standard Deviation |
|---|---|---|---|---|
| BRCA | AE | **0,606** | 0,611 | 0,037 |
| | PreAE | 0,575 | 0,565 | 0,039 |
| | PreAE finetuned | 0,584 | 0,572 | 0,044 |
| KIRC | AE | 0,714 | 0,718 | 0,047 |
| | PreAE | 0,724 | 0,734 | 0,034 |
| | PreAE finetuned | **0,728** | 0,740 | 0,049 |
| UCEC | AE | 0,631 | 0,633 | 0,052 |
| | PreAE | 0,562 | 0,576 | 0,044 |
| | PreAE finetuned | **0,654** | 0,652 | 0,040 |

| | | | | |
|---|---|---|---|---|
| HNSC | AE | 0,573 | 0,571 | 0,034 |
| | PreAE | 0,583 | 0,580 | 0,047 |
| | PreAE finetuned | **0,587** | 0,588 | 0,043 |
| LUAD | AE | **0,641** | 0,653 | 0,065 |
| | PreAE | 0,629 | 0,638 | 0,064 |
| | PreAE finetuned | 0,632 | 0,629 | 0,067 |
| LGG | AE | **0,762** | 0,756 | 0,043 |
| | PreAE | 0,718 | 0,709 | 0,046 |
| | PreAE finetuned | 0,752 | 0,746 | 0,047 |
| LUSC | AE | 0,515 | 0,514 | 0,042 |
| | PreAE | **0,528** | 0,534 | 0,031 |
| | PreAE finetuned | 0,524 | 0,534 | 0,035 |
| COAD | AE | **0,595** | 0,614 | 0,080 |
| | PreAE | 0,531 | 0,556 | 0,071 |
| | PreAE finetuned | 0,570 | 0,548 | 0,065 |
| BLCA | AE | **0,633** | 0,633 | 0,038 |
| | PreAE | 0,609 | 0,625 | 0,041 |
| | PreAE finetuned | 0,622 | 0,623 | 0,036 |
| SKCM | AE | 0,623 | 0,623 | 0,045 |
| | PreAE | **0,633** | 0,628 | 0,038 |
| | PreAE finetuned | 0,626 | 0,610 | 0,044 |
| STAD | AE | **0,614** | 0,631 | 0,050 |
| | PreAE | 0,541 | 0,537 | 0,055 |
| | PreAE finetuned | 0,544 | 0,525 | 0,058 |

**Table S8** Test sets overall spearman correlation statistics for the gene essentiality task for pretrained models. Best performance based on mean results are shown in boldface.

| Representation Model | Mean Metric | Median Metric | Standard Deviation |
|---|---|---|---|
| AE | **0,8554** | 0,8558 | 0,0038 |
| PreAE | 0,8539 | 0,8541 | 0,0030 |
| PreAE finetuned | 0,8551 | 0,8552 | 0,0038 |

| | | | |
|---|---|---|---|
| Exp-DeepDEP (Task-tuned) | 0,8406 | 0,8410 | 0,0035 |
| Exp-DeepDEP (Pretrained) | 0,8335 | 0,8322 | 0,0063 |

**Table S9** Test sets per-gene spearman correlation statistics for the gene essentiality task for pretrained models. Best performance based on mean results are shown in boldface.

| Representation Model | Mean Metric | Median Metric | Standard Deviation |
|---|---|---|---|
| AE | **0,249** | 0,247 | 0,008 |
| PreAE | 0,237 | 0,237 | 0,007 |
| PreAE finetuned | 0,248 | 0,247 | 0,006 |
| Exp-DeepDEP (Task-tuned) | 0,179 | 0,180 | 0,018 |
| Exp-DeepDEP (Pretrained) | 0,067 | 0,042 | 0,108 |

### Supplemental Figures

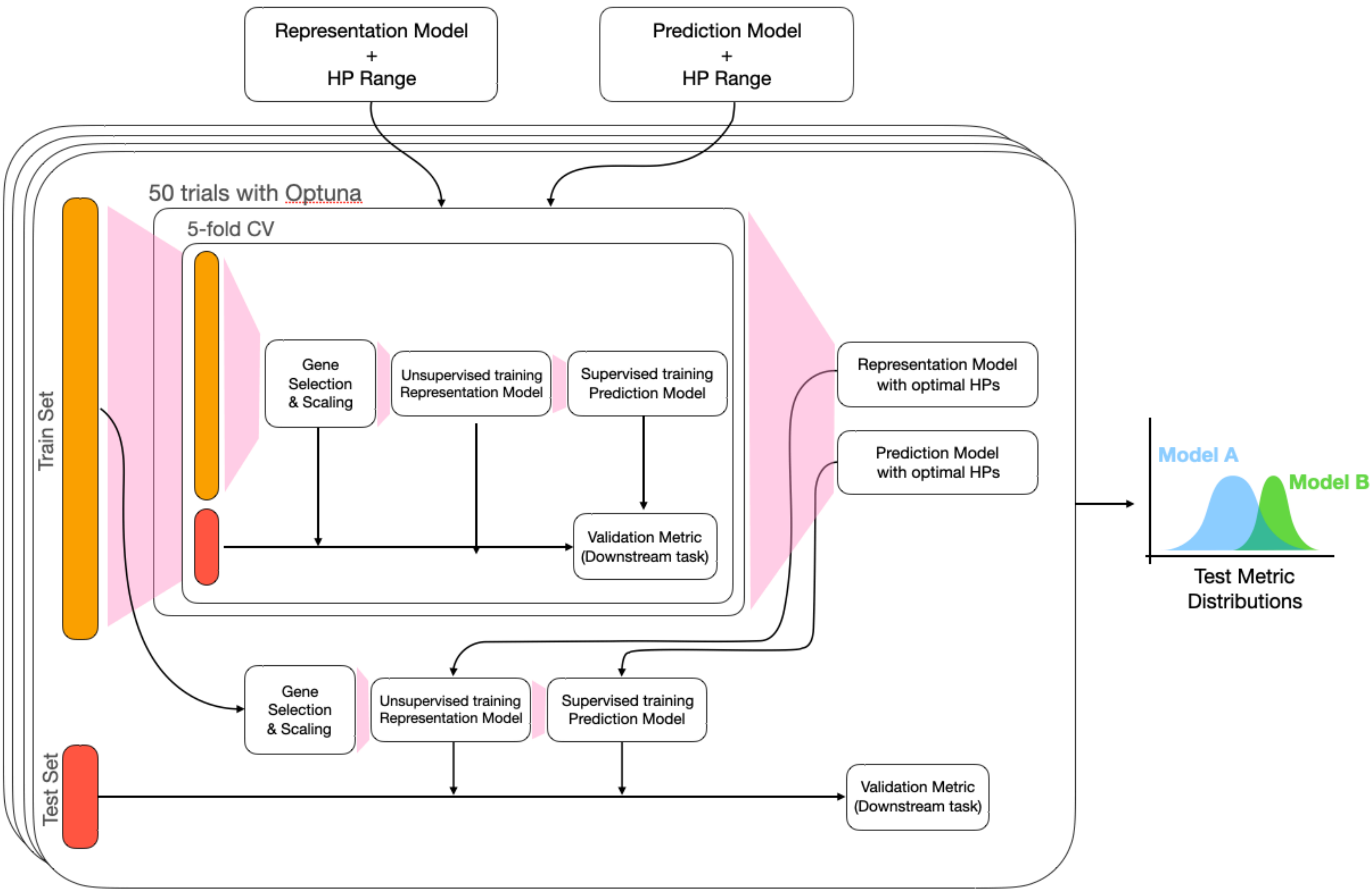


**Supp. Figure S1. Repeated hold-out pipeline description**. The different planes correspond to repetitions of the process, allowing to create the scores distributions per model seen on the right of the figure.

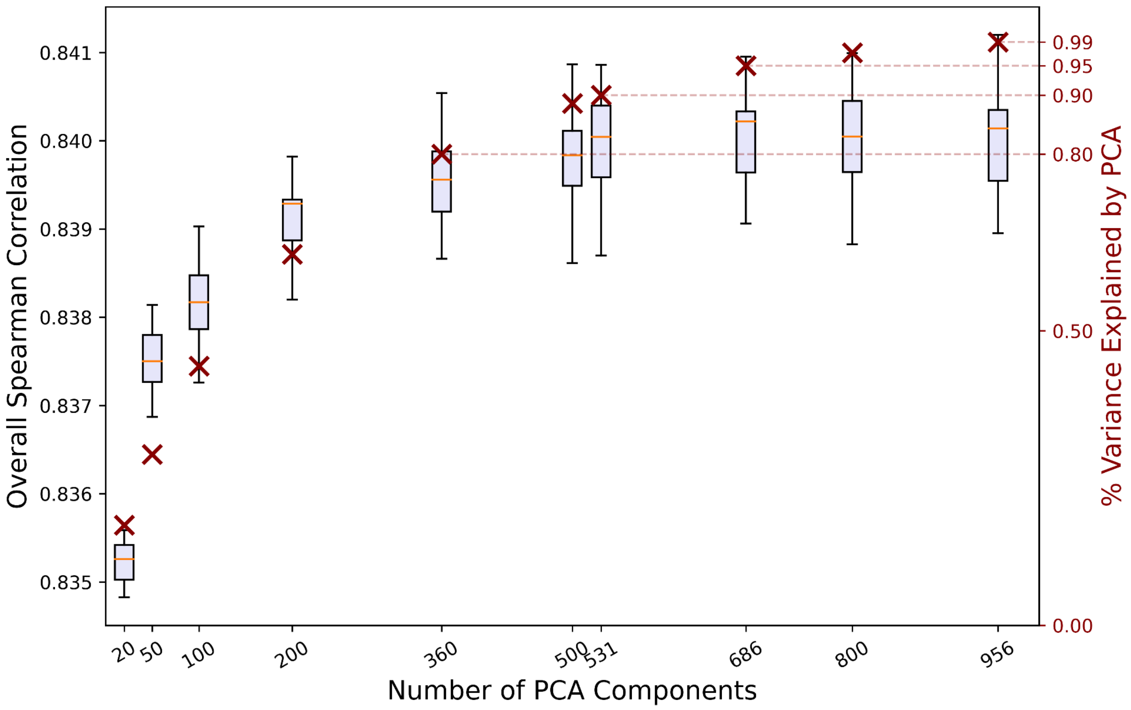


**Supp Figure S2.** Overall spearman correlation for different number of components for the PCA representation of gene fingerprints. Red crosses indicate the percentage of variance explained by PCA for the corresponding number of PCA components. For each tested number of PCA components, the overall spearman correlation is obtained over a 5-fold cross validation run with 12 different hyperparameter sets.

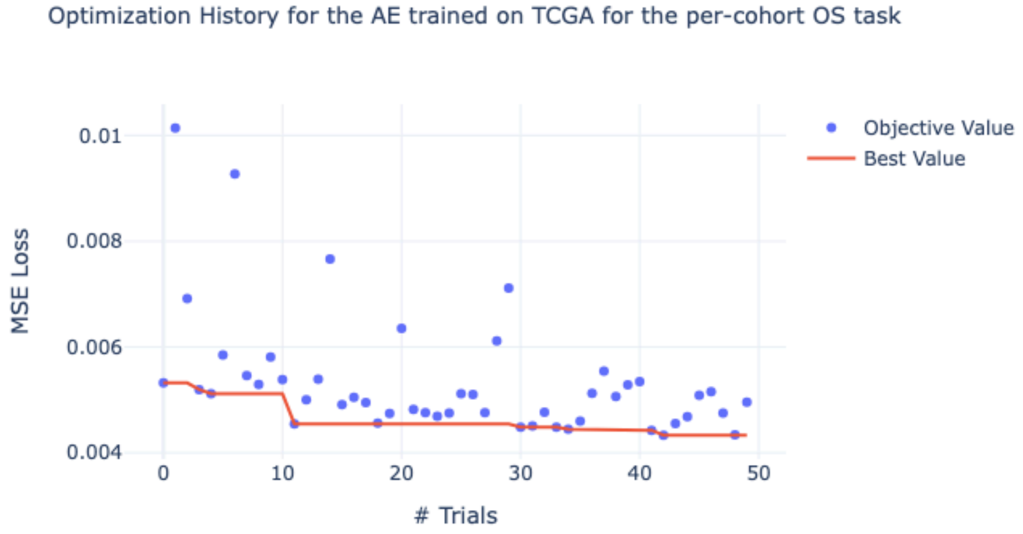


**Supp Figure S3.** Optimization History for the AE trained on 22 cohorts of TCGA excluding cohorts used in the downstream task. After 50 trials the final PreAE for the per-cohort OS prediction task has 255 latent dimensions, no hidden layers, a learning rate of $3.1e-4$ and a dropout rate of $1.1e-2$.

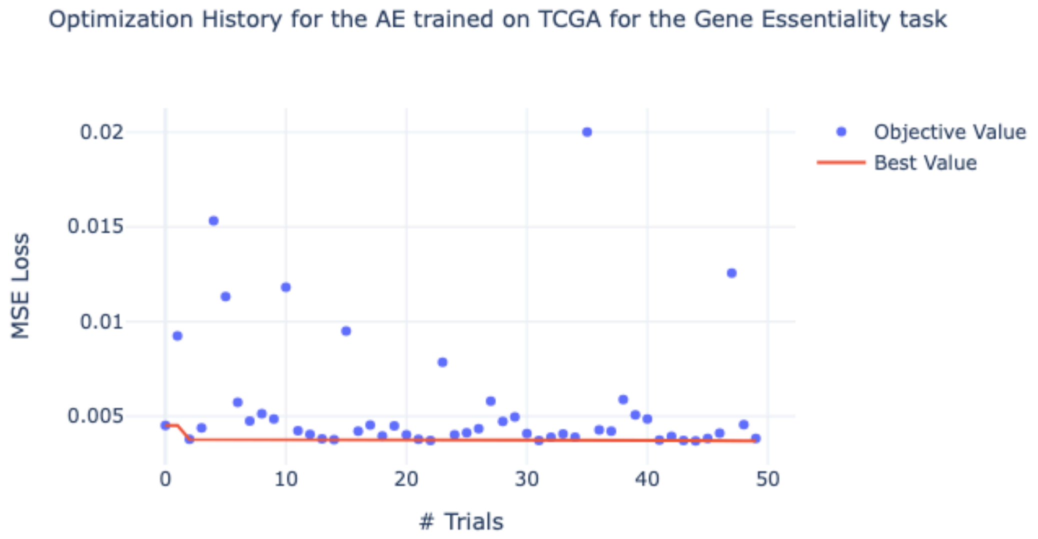


**Supp Figure S4.** Optimization History for the AE trained on the 33 cohorts of TCGA for the Gene Essentiality task. After 50 trials, the final PreAE for the gene essentiality task has 255 latent dimensions, no hidden layers, a learning rate of $1.6e-4$ and a dropout rate of $0.13$.

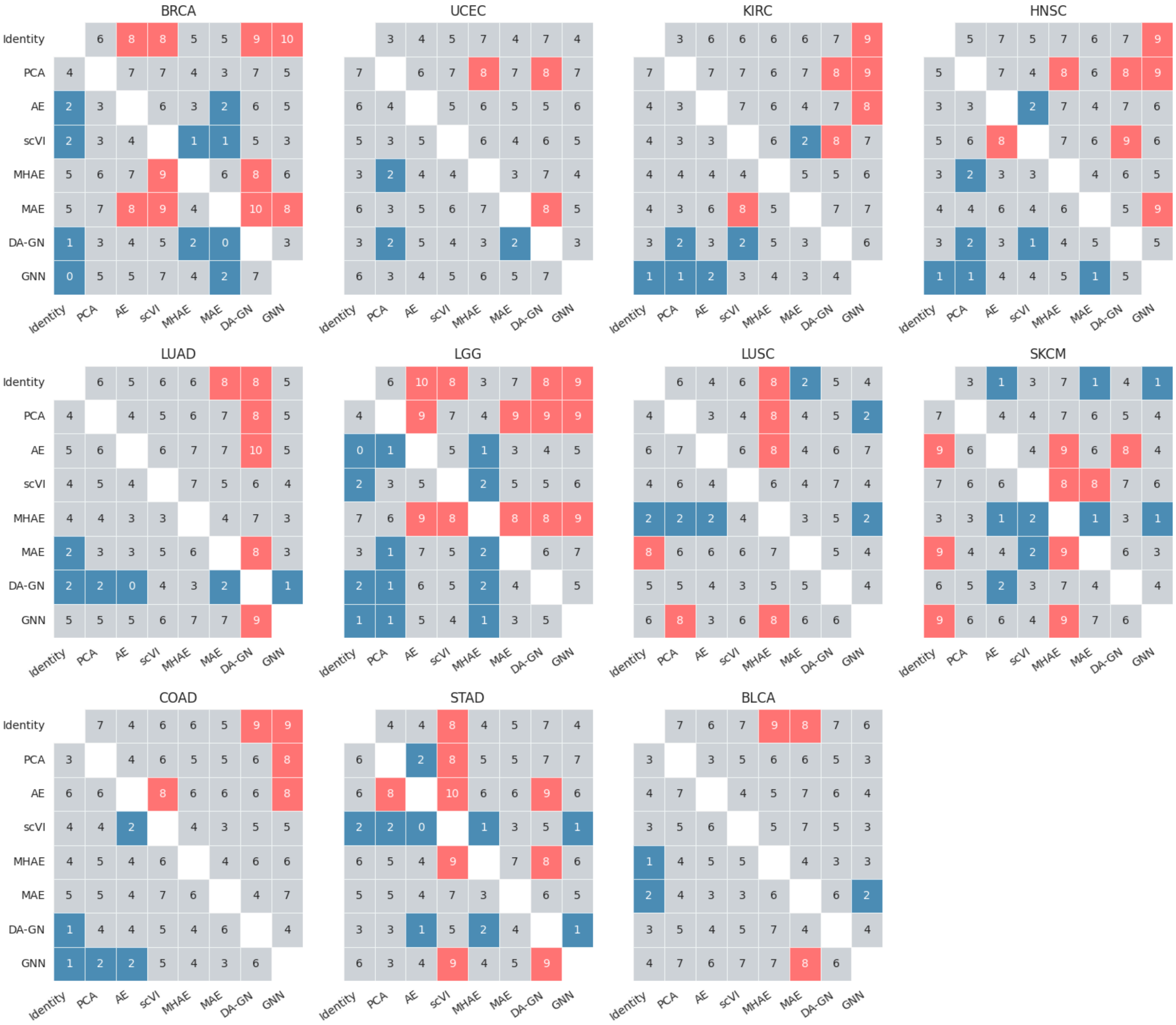


**Supp Figure S5. Comparison of performance on per-cohort OS prediction task on different TCGA cohorts for different bulk RNA-seq representation models.** *Number of folds for which the c-index for the model on the y axis is higher than for the model on the x axis. Red (Blue) boxes indicate 75% acceptance criterion on test folds for superiority is satisfied by the model on the y (x) axis.*

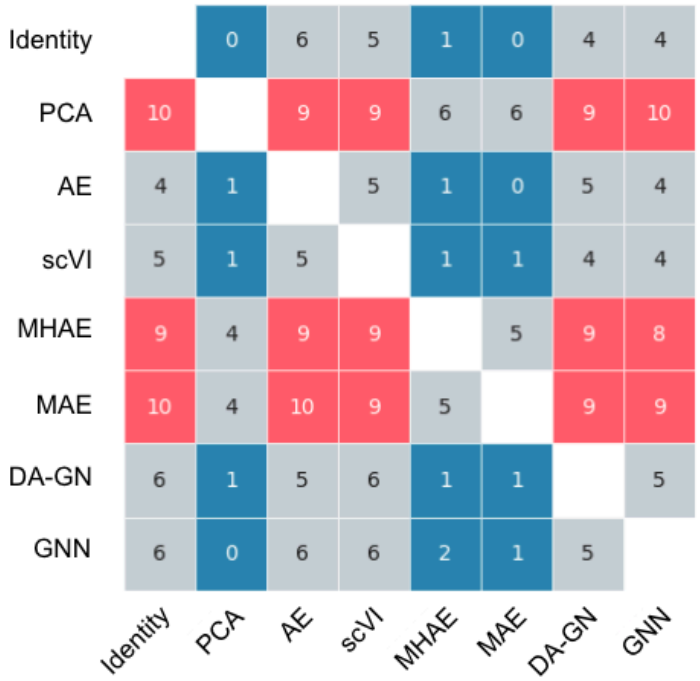


**Supp Figure S6. Comparison of performance on pan-cancer OS prediction task for different bulk RNA-seq representation models.** *Number of folds for which the c-index for the model on the y axis is higher than for the model on the x axis. Red (Blue) boxes indicate 75% acceptance criterion on test folds for superiority is satisfied by the model on the y (x) axis.*

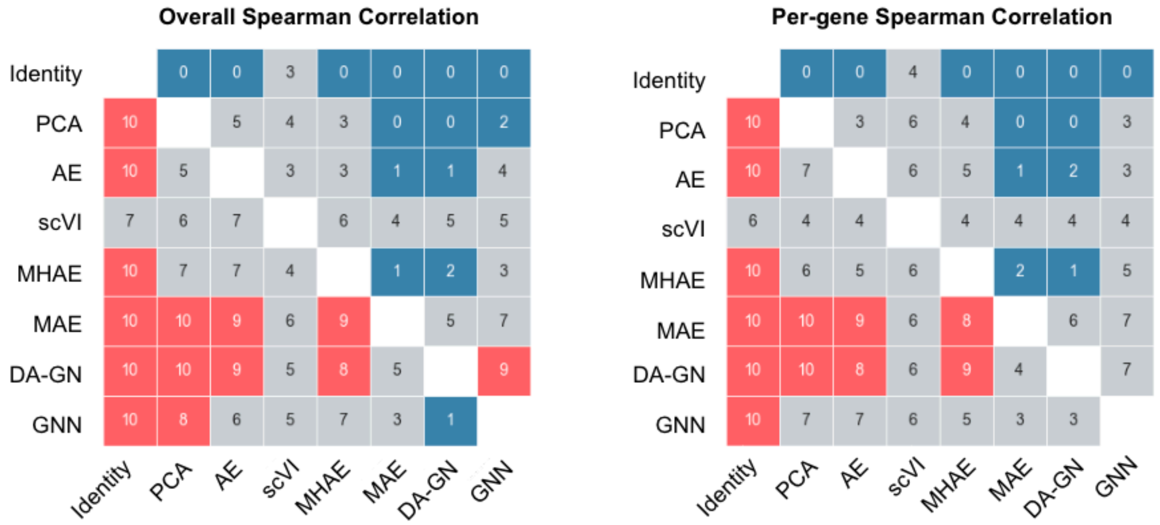


**Supp Figure S7. Comparison of performance on gene essentiality prediction task on DepMap dataset for different bulk RNA-seq representation models.** *Top panel) Number of folds for which the overall correlation for the model on the y axis is higher than for the model on the x axis. Red (Blue) boxes indicate 75% acceptance criterion on test folds for superiority is satisfied by the model on the y (x) axis.* Bottom panel) Same as Top panel, but *correlation computed per-gene.*

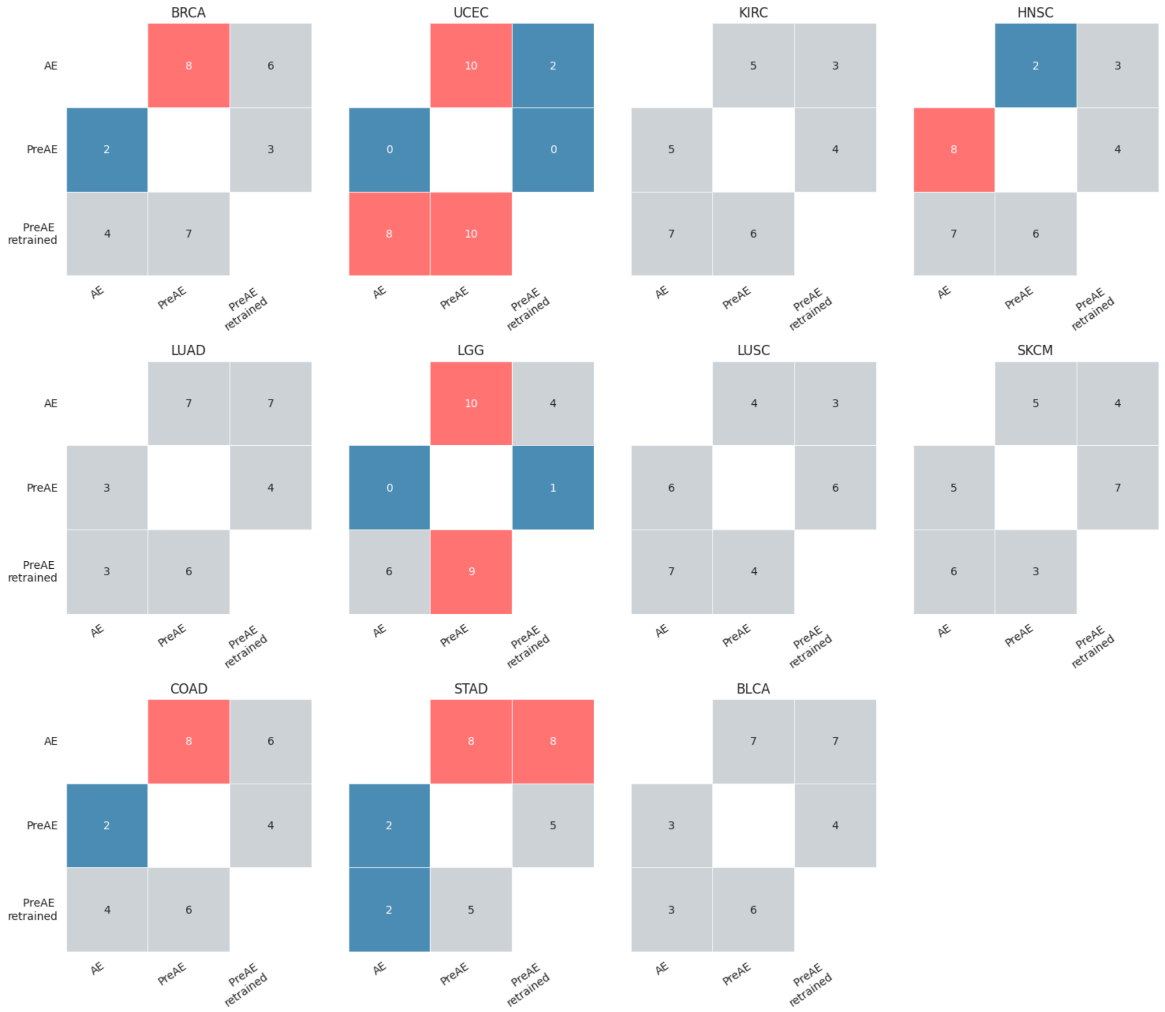


**Supp Figure S8. Comparison of performance on per-cohort OS prediction task on different TCGA cohorts for pretraining experiments.** *Number of folds for which the c-index for the model on the y axis is higher than for the model on the x axis. Red (Blue) boxes indicate 75% acceptance criterion on test folds for superiority is satisfied by the model on the y (x) axis.*

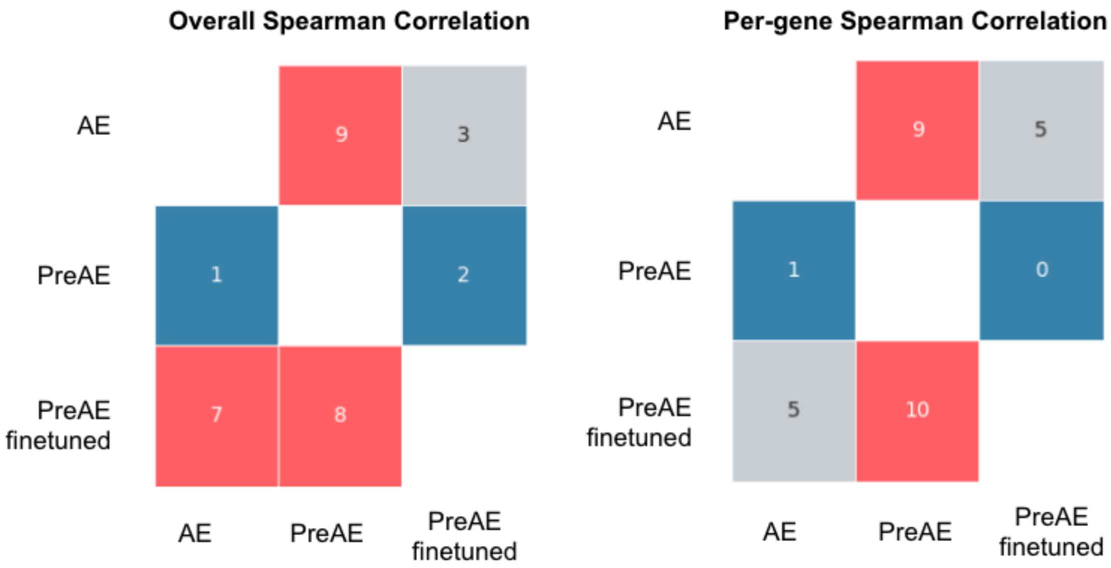


**Supp Figure S9**. **Comparison of performance on gene essentiality prediction task on CCLE dataset for different bulk RNA-seq representation models.** *Left panel) Number of folds for which the overall correlation for the model on the y axis is higher than for the model on the x axis. Red (Blue) boxes indicate 75% acceptance criterion on test folds for superiority is satisfied by the model on the y (x) axis.* *Right panel) Same as Top panel, but correlation computed per-gene.*

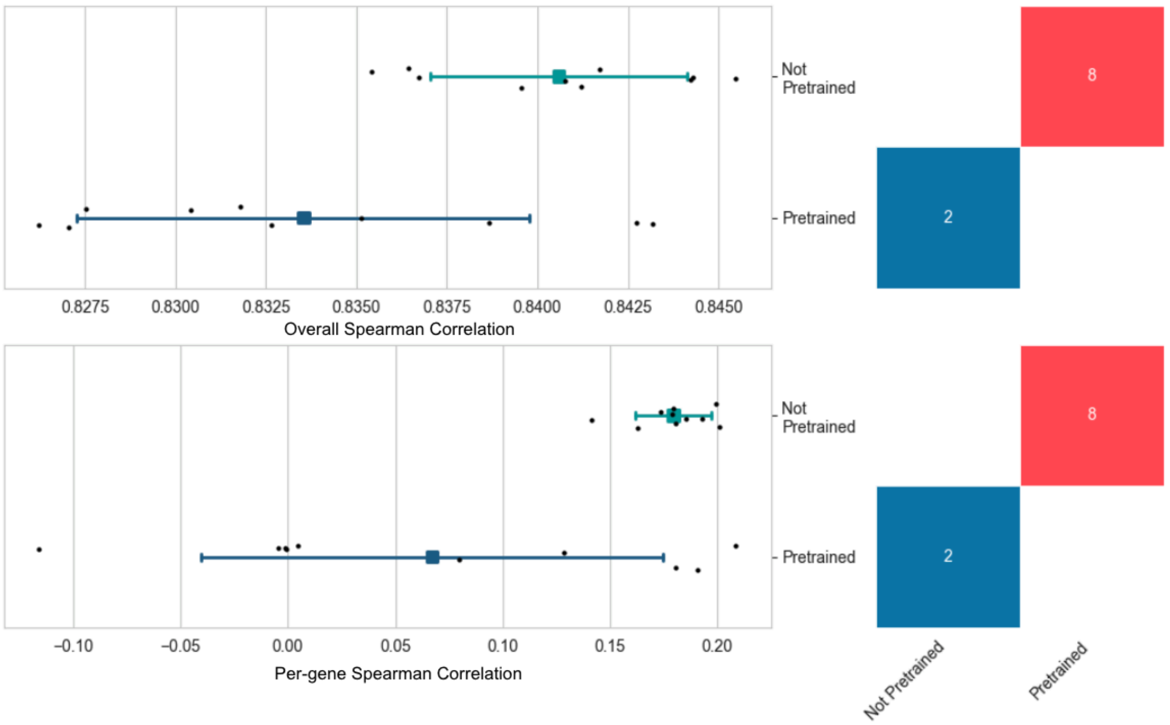


**Supp Figure S10.** Impact of pre-training an Exp-DeepDEP architecture on our data splits using the 5,000 most variable genes from TCGA as features. Skipping the pretraining step of

Exp-DeepDEP seems to help reach better performances both in overall Spearman correlation and per-gene Spearman correlation, contrary to experiments performed on the multimodal DeepDEP model.
